# BtuB TonB-dependent transporters and BtuG surface lipoproteins form stable complexes for vitamin B_12_ uptake in gut *Bacteroides*

**DOI:** 10.1101/2022.11.17.516869

**Authors:** Javier Abellon-Ruiz, Kalyanashis Jana, Augustinas Silale, Andrew M. Frey, Arnaud Baslé, Matthias Trost, Ulrich Kleinekathöfer, Bert van den Berg

## Abstract

Vitamin B_12_ (cobalamin) is the most complex vitamin and essential for many human gut microbes. However, cobalamin is synthesised only by a limited number of bacteria, making many gut microbes dependent on scavenging to meet their cobalamin requirements. Since bacterial densities in the gut are extremely high, competition for cobalamin is severe, making it a keystone micronutrient that shapes human gut microbial communities. Contrasting with Enterobacteria like *Escherichia coli* which only have one outer membrane (OM) transporter dedicated to B_12_ uptake (BtuB), members of the dominant genus *Bacteroides* often encode several vitamin B_12_ OM transporters together with a conserved array of surface-exposed B_12_-binding lipoproteins. Here we show, via X-ray crystallography, cryogenic electron microscopy (cryoEM) and molecular dynamics (MD) simulations, that the BtuB1 and BtuB2 transporters from the prominent human gut bacterium *Bacteroides thetaiotaomicron* form stable complexes with the surface-exposed lipoproteins BtuG1 and BtuG2. The lipoproteins cap the external surface of their cognate BtuB transporter and, when open, capture B_12_ via electrostatic attraction. After B_12_ capture, the BtuG lid closes, with concomitant transfer of the vitamin to the BtuB transporter and subsequent transport. We propose that TonB-dependent, lipoprotein-assisted small molecule uptake is a general feature of *Bacteroides spp*. that is important for the success of this genus in colonising the human gut.

## INTRODUCTION

Vitamin B_12_ (cobalamin) is a complex organometallic cofactor and the most complex vitamin^1^, consisting of a corrin ring containing a cobalt atom in the centre coordinated with an upper ligand (such as adenosyl or methyl group) and a lower ligand anchored to the ring through a nucleotide loop (Fig. 1a)^2^. The upper ligand contains the chemical reactivity, directly participating in reactions, while the lower ligand provides functional specificity^3–5^. There are three different families of lower ligands; benzimidazoles, purines and phenolics^5^. The nature of this ligand is important as enzymes belonging to different species can require different lower ligands to be active^3,4,6^. Vitamin B_12_ is involved in a wide variety of metabolic processes, and for many organisms it is an essential cofactor for the final enzymatic reaction of the L-methionine biosynthesis pathway in the cytoplasm^7,8^. Despite its many roles in eukaryotic and prokaryotic cells, only a small group of microorganisms is able to produce vitamin B_12_. However, its synthesis is energetically expensive, requiring approximately 30 enzymatic steps^9^. As a result, microorganisms have developed mechanisms to take up exogenous cobalamins, named the salvage route. In Gram-negative bacteria, the translocation of cobalamins presents challenges given that three different compartments need to be crossed: the outer membrane (OM), the periplasm and the inner membrane (IM). The best-characterised B_12_ transport system in Gram-negatives is that of *Escherichia coli*, comprising the OM TonB-dependent transporter (TBDT) BtuB, the periplasmic binding protein BtuF, and the BtuCD ABC transporter located in the IM^10–12^.

**Fig. 1.**
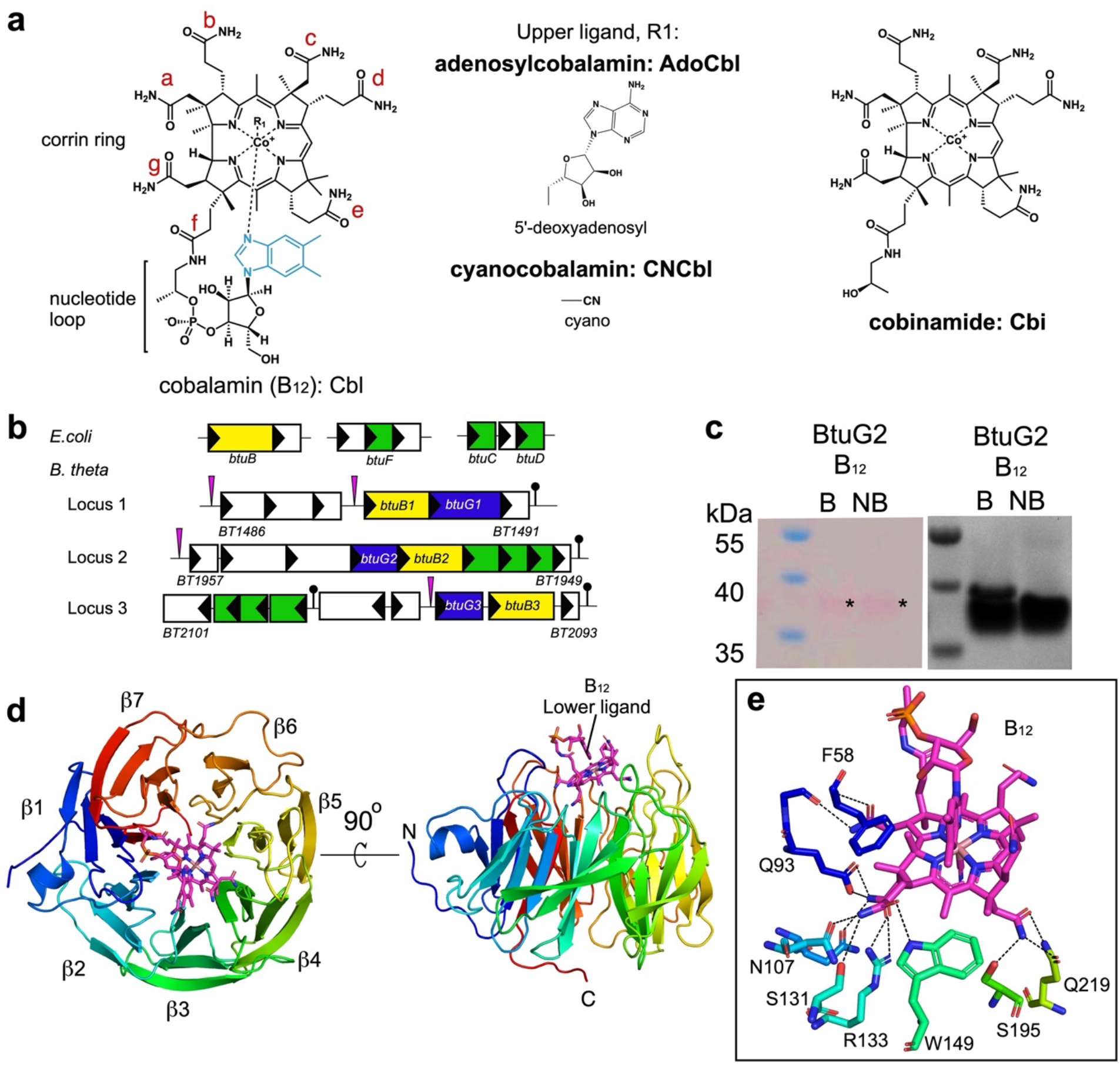
BtuG2 has a beta propeller fold which binds corrinoids. **a**, Diagrams showing the structures of the corrinoids used for crystallography. Lower ligand is depicted in light blue in the “base-on” conformation. Side chains of the corrin ring are labelled with letters in red. **b**, Genetic organization of the B_12_ transport system in *E. coli* (*btuBFCD*) and the three homologous loci in *B. theta*, showing the locations of BtuB (yellow) and BtuG proteins (blue). The pink triangles represent B_12_ dependent riboswitches and the black lollipops transcription terminators. The inner membrane ABC transporters are in green. **c**, SDS gel showing faint pink bands, indicated with an asterisk on the left panel, corresponding to CNCbl. The boiled sample also shows a minor lower mobility band (∼15% of the sample) which corresponds to the fraction of BtuG2 that has lost CNCbl after boiling. The right panel shows the same gel after Coomassie staining (B, boiled; NB, non-boiled). **d**, Cartoon representation of BtuG2 (in rainbow colour, N terminus in blue) bound to CNCbl (magenta). Note that the lower ligand is pointing outwards (right panel). **e**, Close-up of the residues forming hydrogen bonds (black dashed lines) with CNCbl.

The human gut microbiome is the highly complex community of microorganisms in the gastrointestinal tract and has been implicated in many aspects of human health^13,14^. The vast majority of human commensal microbes is located in the distal gut with bacterial densities of ∼10^12^/g luminal content^15^, making competition for resources likely severe. To understand the requirements for being competitive within the gut it is important to understand how small-molecule acquisition occurs. Despite this, most of these uptake processes are poorly characterised, especially for non-Proteobacterial members of the gut microbiota.

The Gram-negative bacterium *Bacteroides thetaiotamicron* (*B. theta*), like the vast majority of the Bacteroidetes phylum, cannot make vitamin B_12_ but possesses several B_12_-dependent enzymes^16^. One of these enzymes is the methionine synthase MetH, making B_12_ an essential nutrient for this bacterium. *B. theta* therefore serves as a model for the salvage routes of cobalamins by the most abundant Gram-negative phylum in the gut^17^. Interestingly, while model organisms like *E. coli* have just one vitamin B_12_ uptake locus, *B. theta* has three, together containing 24 proteins (Fig. 1b), suggestive of the importance of B_12_ uptake for this organism^18^. All three loci contain one copy each of the TonB dependent transporter (TBDT) BtuB (BtuB1-3) as well as one copy of BtuG, which is always located adjacent to BtuB on the genome^19^. The presence of multiple outer membrane B_12_ transporters is a widespread feature in *Bacteroides* (and Bacteroidetes in general), with up to four BtuBs^18^. Competition assays have shown that strains lacking *btuB2* or *btuG2* have fitness defects *in vivo* and *in vitro*, and are rapidly outcompeted by wild type cells, suggesting that locus 2 might encode the primary uptake system in *B. theta*^18^. Interestingly, BtuG2 is a surface-exposed OM lipoprotein with an extremely high affinity for B_12_ (sub-pM) and is able to remove the vitamin from human intrinsic factor (IF), which binds B_12_ in the small intestine prior to absorption^19^. However, while BtuG2 is probably highly efficient in scavenging B_12_ from the gut lumen, it is unclear how the vitamin is transferred to BtuB2 for uptake. Genomic studies show that all *Bacteroidetes* B_12_ transport loci include homologs of BtuB and almost all encode BtuG, suggesting the two proteins work together to bring B_12_ into the cell^18,19^.

Here we present X-ray crystal structures of *B. theta* BtuG2 complexed with cyanocobalamin (the manufactured and most stable form of B_12_, CNCbl), adenosylcobalamin (one of the biologically active forms of vitamin B12 with the bulkiest upper ligand, AdoCbl) as well as the precursor cobinamide (Cbi), which lacks the lower ligand and the nucleotide loop. The positively charged cofactors (Co is charged +3)^20^ are bound at the same site on the negatively charged face of the BtuG2 beta-propeller and make many polar and hydrophobic interactions with protein residues. Molecular dynamics (MD) simulations reveal that BtuG2 can attract cyanocobalamin to its binding site over large distances (> 35 Å), establishing it as an efficient B_12_ capturing device. BtuG2 forms a stable complex with BtuB2 in the OM, allowing co-purification and structure determination of BtuB2G2 by X-ray crystallography. The structure of the BtuB2G2 complex demonstrates that BtuG2 caps the transporter, reminiscent to recently described SusCD systems involved in glycan uptake^21^. Interestingly, however, the BtuB2G2 structure as well as the cryo-EM structure of the BtuBG complex from locus 1 (BtuB1G1) show closed transporters in the absence of substrate, which is different from SusCD systems. Steered and conventional MD simulations suggest a pedal bin uptake mechanism and provide clues about how B_12_ is transferred from BtuG2 to BtuB2 for subsequent uptake. Together with OM proteomics data, our study suggests that lipoprotein-assisted small molecule uptake operates for most, and perhaps all, TBDTs of Bacteroides spp, and we propose that this is one reason why these microbes are so successful within the human gut.

## RESULTS

### BtuG2 binds a variety of corrinoids

To identify the B_12_ binding site of BtuG2, we overexpressed the protein without its signal sequence and lipid anchor in *E. coli*. To load BtuG2 with the ligand before the size exclusion chromatography (SEC) purification step, we added CNCbl at a 1:2 molar ratio (protein:vitamin). The non-bound vitamin elutes later than the BtuB2-CNCbl complex, allowing us to eliminate excess vitamin in the sample. As observed previously, CNCbl binds to BtuG2 with very high affinity^19^. The BtuG2-CNCbl complex is extremely stable, given that a boiled sample run on a denaturing SDS-PAGE gel shows a visible pink band around 40 kDa, suggesting that some CNCbl is still bound to BtuG2 (Fig. 1c). An initial BtuG2-CNCbl structure was solved to 1.9 Å resolution using cobalt-based single-wavelength anomalous diffraction (Co-SAD), showing unambiguous electron density for CNCbl. This model was used to solve an additional dataset at slightly higher resolution (1.7 Å) (Supplementary Table 1). BtuG2 adopts a seven-bladed beta-propeller fold (Fig. 1). All blades have 4 beta strands except blade number 6 (β6), which has only two. The propeller blades define a central cavity which forms the B_12_ binding pocket with an interface area of 698 Å^2^ as measured via PISA^22^. CNCbl is oriented with the cyano group (upper ligand) pointing towards the protein centre while the lower ligand, present in its “base-on” conformation^20^, is completely exposed to the exterior, with the nucleotide loop sitting in the cleft formed between blades 1 and 7 (Fig. 1). There are 13 hydrogen bonds between BtuG2 and CNCbl, all mediated by the amide groups of the substituents from side chains a, b, c and g of the corrin ring (Supplementary Fig. 1b and Fig. 1d) and involving protein residues from blades 1 to 4. CNCbl binding is further stabilised by Van der Waals forces provided by 11 residues, evenly distributed across the blades. The cavity that contains the cyano upper ligand is narrowed by Trp194, Trp272 and Tyr316 (numbering based on the annotated Uniprot sequence, including the signal peptide). To explore if this narrow pocket could accommodate a cobalamin with a bulky upper ligand, we obtained a structure to 2.3 Å resolution of BtuG2 co-crystallised with AdoCbl. In this structure the side chain of Trp272 has rotated, enlarging the cavity to fit the adenosyl group. The interaction is stabilised by stacking forces from Trp272 and a hydrogen bond between the hydroxyl group of Tyr335 and the nitrogen in position 3 of the pyrimidine ring of the adenosyl group (Supplementary Fig. 1c). For both structures the hydrogen bonds interacting with the side chains of the corrin ring are identical. A Consurf analysis^23^ shows that while the degree of conservation of the protein surface is low overall, the residues involved in forming hydrogen bonds with the ligands are highly conserved (Supplementary Fig. 2). Thus, given that the lower ligand does not interact with the protein, the structures suggest that BtuG2 can bind many different, if not all, corrinoids. To investigate whether the lack of lower ligand would affect the binding mode, we also determined the BtuG2-Cbi crystal structure, using data to 1.35 Å resolution. The orientation of the ligand is virtually the same as for the other structures (Supplementary Fig. 1d), confirming BtuG2 as a versatile corrinoid binder.

### Long-range electrostatic attraction of CNCbl by BtuG2

To study the BtuG2-CNCbl interaction in real time, unbiased MD simulations were performed using the crystallographic model for CNCbl-BtuG2. For analysis, we defined the B_12_ binding pocket as those residues that have an atom within 3 Å distance of the ligand (Supplementary Fig. 3). During the 1 µs-long simulation, the centre of mass (COM) distance between the ligand and the binding pocket varied less than 1 Å, indicating that CNCbl is bound very strongly and only wiggles slightly within the binding pocket (Fig. 2a). The number of hydrogen bonds fluctuates around a constant number (Supplementary Fig. 3) and a superposition of the ligand before and after the simulation shows an almost identical orientation, supporting the notion that the interaction is highly stable (Supplementary Fig. 3). A calculation of the short-range electrostatic interactions between the ligand and the binding pocket results in large electrostatic interaction energies of around -30 kcal/mol throughout the simulations, explaining the stability of the ligand within the pocket.

**Fig. 2.**
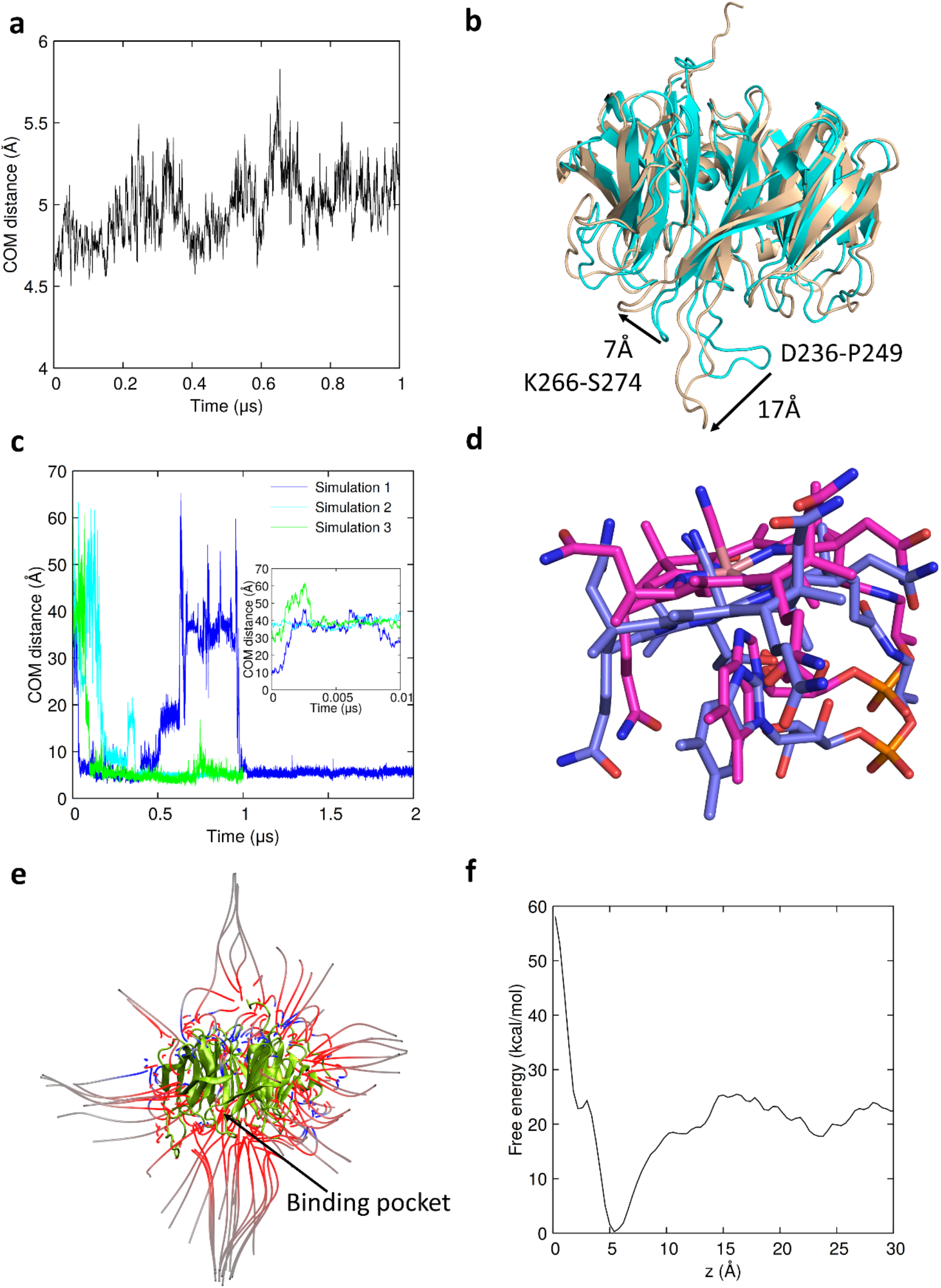
BtuG2-CNCbl binding dynamics. **a**, COM distance between the BtuG2 binding pocket and CNCbl for simulations starting from the crystal structure. **b**, Superposition of relaxed BtuG2 and the crystal structure. The displaced loops are annotated (cyan for the crystal structure, wheat for the relaxed structure). Arrows indicate loop movements away from the binding pocket during relaxation **c**, COM distances between the binding pocket of the relaxed structure and the ligand initially placed arbitrarily at 8.5, 37.1 and 34.9 Å. The unbiased simulations 2 and 3 where stopped after 1µ as CNCbl was stably bound. **d**, Representative overlay of the final ligand position at the end of the unbiased simulation 1 (carbon colour blue) and the position in the crystal structure (carbon colour magenta). **e**, Representation of the electric field of BtuG. A large number of electric field lines originate from the binding site of BtuG, caused by an accumulation of negative charges in the protein. The neutral CNCbl has a large dipole moment and is attracted by the binding site. The red field lines point towards the negative charges of the protein and the blue ones towards the positive charges. Line density indicates the strength of the electric field. **f**, Free energy profile between CNCbl and BtuG2 as a function of the COM distance between the two binding partners. The free energy of binding, i.e., the difference between the lowest energy and the energy at large COM distances, is about 20 kcal/mol.

In order to assess the CNCbl binding mechanism we analysed apo-BtuG2 using unbiased MD simulations. To this end, the CNCbl molecule was removed from the crystallographic model and the structure was simulated for 1 µs. The structure relaxed during the first half of the simulation and did not significantly change thereafter. In the absence of the ligand, loops Asp236-Pro249 and Lys266-Ser274 (both from blade 5, henceforth named β5A and β5B loops) moved outwards away from the binding site by 17 and 7 Å, respectively (Fig. 2b). It is worth pointing out that loop β5A contains Tyr239 which, in the crystal structures, contributes to CNCbl and AdoCbl binding through van der Waals (vdW) forces or via a hydrogen bond with Cbi. Additionally, loop β5B contains Trp272, providing vdW forces for the CNCbl and Cbi binding. Moreover, the side chain of this residue rotates to allow AdoCbl binding (Supplementary Fig. 1). No significant structural changes other than these two displaced loops and the C terminus were detected upon removal of CNCbl.

To explore the efficiency of CNCbl capture, we next ran three unbiased MD simulations of 4μs in total, each starting with a CNCbl molecule placed at a different arbitrary initial position 8.5, 37.1 and 34.9 Å away from the binding pocket (Supplementary Fig. 4). Strikingly, in all these simulations, regardless of the initial distance and orientation of the CNCbl molecule, the ligand moved rapidly into the binding site. The COM distance analysis shows that during the first µs, CNCbl binds via an association-dissociation process during which the ligand can move away from BtuG2 (sometimes almost doubling the furthest starting point distance and reaching the maximum distance possible in the present periodic simulation box) (Supplementary movie 1). Once the ligand is bound in the right conformation, it stays in the pocket for the remainder of the simulation (Fig. 2c). This strong attraction over long distances suggests a long-range electrostatic interaction. In all three simulations but after different times, CNCbl eventually adopts a similar conformation to the crystal structure (Fig. 2d). BtuG2 has many negative charges (net charge -18 e) leading to a strong electric field that attracts CNCbl to the binding pocket. A dense set of electric field lines points towards the binding pocket, explaining the attraction of an electrically neutral but strongly dipolar ligand with a dipole moment of 6.7D towards the binding pocket (Fig. 2e and Supplementary Figs. 4a and 5a,b). Besides the electrostatic interactions, our MD studies show that conformational changes from loop β5A play a key role in ligand binding. Due to the displacement of this loop when relaxing the apo-BtuG structure, the binding pocket became much bigger than in the crystal structure. Upon binding, the ligand is held by loops β5A and Asn54-Thr64 like a pair of pliers (Supplementary Fig. 3 and Supplementary movie 1). Free energy calculations show that the binding free energy for CNCbl is around ΔG=20.0 kcal/mol (Fig. 2f). This corresponds to a dissociation constant of 1.4 × 10^−14^ M calculated using *k* = *e*^−Δ*G*/*RT*^ at 300 K, which is in reasonable agreement with the experimental value of 1.93 × 10^−13^ M reported previously^19^, considering the calculation approximations involved and the difficulties in measuring extreme affinities via surface plasmon resonance.

To better understand ligand binding, we calculated the short-range electrostatic interaction energy between the highly dipolar CNCbl and the negatively charged binding site of BtuG2 along the unbiased MD trajectories. Comparing the configurations and electrostatic energies suggests that binding occurs for values below -20 kcal/mol. Thus, once the electrostatic interaction energy between CNCbl and BtuG2 reaches values below -20 kcal/mol, CNCbl is strongly bound to the binding site of BtuG2 (Supplementary Fig. 4c), as supported by the number of hydrogen bonds formed during the simulations (Supplementary Fig. 4d).

### BtuB2 forms a stable complex with BtuG2

The presence of *btuB2* in an operon with *btuG2* suggests that the proteins could be working together in B_12_ acquisition. To explore this possibility, we added a C-terminal hexa-histidine tag to chromosomal *btuB2* and replaced the wild type promoter with a strong constitutive promoter (Methods). Expression levels of BtuB2 in minimal media containing methionine were low (∼0.05-0.1 mg/l culture), but sufficient for purification via IMAC and SEC. Analysis of the sample on a SDS page gel showed two clear bands upon boiling the sample (75 kDa and 40 kDa) (Fig. 3a). The molecular weight of the upper band agrees with that of BtuB2 (77 kDa; Bt1953) and mass spectrometry analysis demonstrated that the ∼40 kDa band corresponds to BtuG2 (Bt1954). Without boiling only one band is present, indicating that both proteins form a complex even in SDS.

**Fig. 3.**
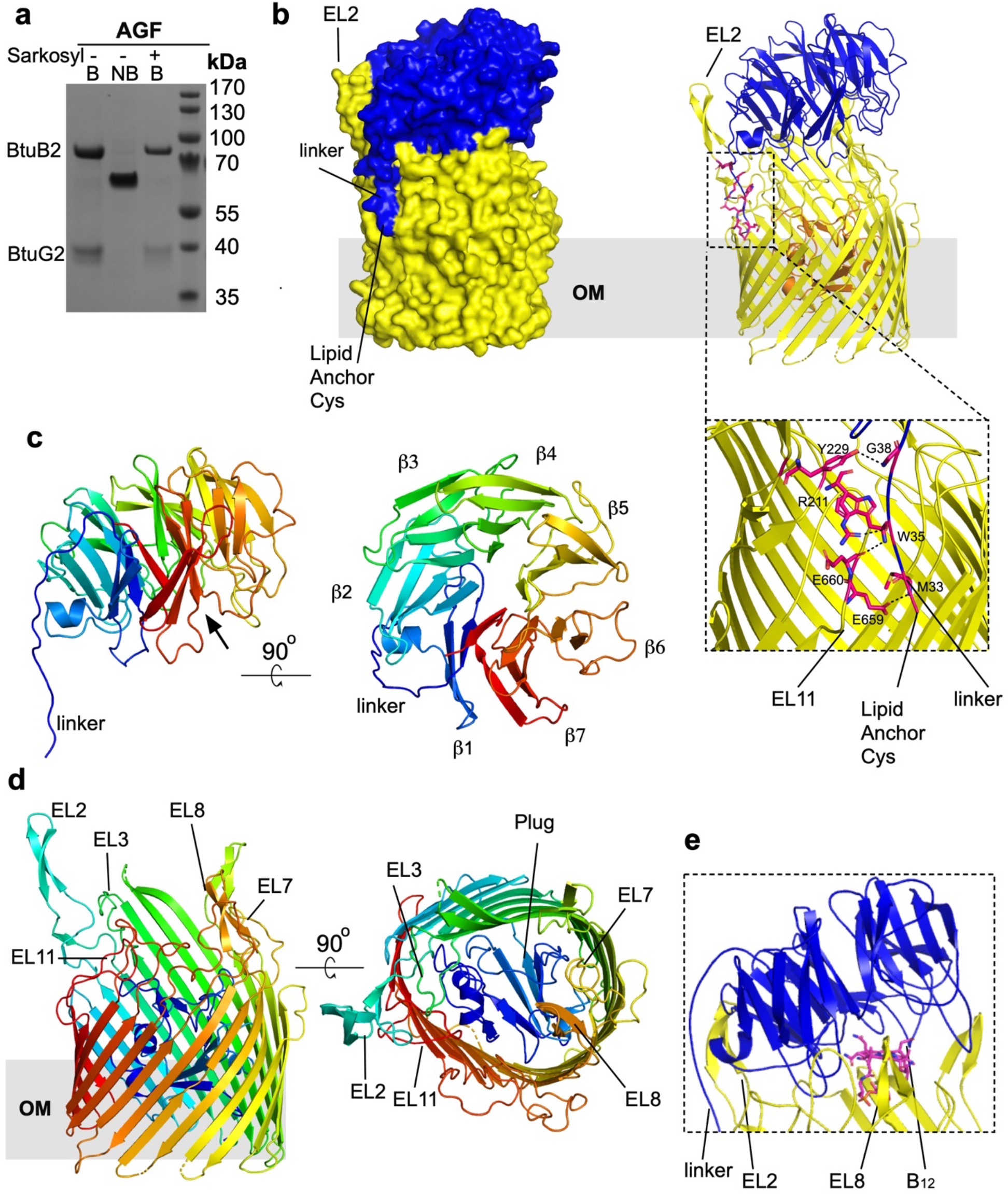
BtuG2 and BtuB2 form a stable complex. **a**, Gel showing that BtuG2 co-purifies with BtuB2. B, boiled samples, NB non-boiled sample. A sarkosyl pre-extraction step did not improve the purity of the sample and suggests that the ∼40 kDa band is an OMP. **b**, Surface (left panel) and cartoon (right panel) views from the side for BtuB2G2 (BtuB2 β-barrel in yellow, plug in orange; BtuG2 in blue). EL denotes extracellular loop, β denotes β-propeller blade. The close-up view shows hydrogen bonds stabilising the linker (black dashed lines, residues in red sticks). For clarity, loops are smoothed and the plug has been removed in this close-up view. **c**, Lateral view of BtuG2 in rainbow colouring (orientation as in b; N-terminus in blue) and rotated by 90° to show the exterior surface. Black arrow indicates the B_12_ binding site. **d**, Lateral view of BtuB2, and rotated 90° showing the plugged pore. **e**, Close-up view of the cartoon representation of BtuG2-BtuB2 (smooth loops) showing the BtuG2 CNCbl binding site in the complex. View was obtained after superimposing the BtuG2-CNCbl crystal structure onto BtuG2 in the complex). Note how EL8 from BtuB2 would clash with CNCbl.

The structure of the complex (BtuB2G2) was solved to 3.7 Å resolution by Molecular Replacement (Supplementary Table 1). The structure shows a typical TBDT fold for BtuB2 with an N-Terminal plug domain (pfam Domain PF07715) occluding a 22-stranded-β-barrel (pfam Domain PF00593). There is no electron density for the TonB box, suggesting that the N-terminal region is disordered despite the absence of ligand. BtuG2 is located on top of the barrel, forming an extracellular lid with a large buried interface area of 3,463 Å (Fig. 3b). The BtuG2 model is complete, and shows the lipid anchor of the N-terminal cysteine (residue 32 in the full-length sequence) at the back of the complex (Fig. 3b,c). The linker between Cys32 and the beta propeller runs parallel to extracellular loop 2 (EL2; Ser202-Tyr239) from BtuB2 (Fig. 3b,d) and is constrained by the BtuB2 barrel via 4 hydrogen bonds provided by EL2 and EL11 (Fig. 3b and Supplementary Fig. 6) (EL: <u>e</u>xtracellular loop). The BtuG2-BtuB2 interface is stabilised by ∼40 hydrogen bonds that are evenly distributed across the interface, and 6 salt bridges. (Fig. 3 and Supplementary Fig. 6). The electrostatic interactions between the BtuG2 linker and EL2/EL11 offer a stable anchoring point which could be acting as a hinge to allow the opening of the lid. Gratifyingly, based on the BtuG2-CNCbl structure, the B_12_ binding site of BtuG2 faces BtuB2. However, the BtuG2 lid caps BtuB2, and the BtuG2 binding site is solvent excluded and clearly inaccessible to any B_12_. Moreover, despite the presence of 1.5 molar equivalents of CNCbl during crystallisation, no ligand is present in the crystallised complex. Indeed, BtuB2 loop EL8 is occupying the B_12_ binding pocket region (Fig. 3e) and would clash with the ligand. Therefore, for BtuG2 to be able to bind B_12_, a conformational change is required that opens the BtuG2 lid to expose the binding site. However, it was not possible to load purified BtuB2G2 with CNCbl *in vitro* without destabilising the complex, suggesting that the closed structure is very stable. A ConSurf analysis^23^ shows that the TBDT surface and periplasmic loops are poorly conserved, unlike the BtuB2 extracellular cavity which is highly conserved and contains the B_12_ binding site. (Supplementary Figs. 2 and 7)

### The cryo-EM structure of BtuB1G1 also shows a closed complex

Previous work on SusCD complexes and the peptide transporter RagAB has shown that X-ray crystallography selects for closed states of dynamic transporters^21,24^. To obtain more experimental insight into the dynamics of BtuG lid opening, we determined the structure of the BtuB1G1 (*i*.*e*. the BtuBG complex of locus 1; Fig. 1b) complex by cryo-EM. BtuG1 is ∼70 kDa in size due to the presence of a C-terminal extension of ∼250 residues that is homologous to the BtuH2 protein that was recently shown to bind B_12_ on the cell surface of *B. theta*^25^; thus, BtuG1 contains two B_12_ binding sites The additional BtuH domain increases the size of the extra-membrane part of the complex substantially, facilitating structure determination by cryo-EM. We obtained ∼3.2 Å maps for the complex in the absence of CNCbl (Fig. 4a) (Supplementary Table 2). The additional BtuH domain (BtuG1 H domain) is clearly visible and appears to be docked to BtuG1, in a position that would not allow B_12_ binding (Fig. 4d). The resolution of the BtuH domain was substantially worse than the rest of the complex, implying some degree of conformational flexibility. Interestingly, and contrasting with substrate-free structures of SusCD glycan transporters, the BtuB1G1 EM structure is very similar to the BtuB2G2 crystal structure with a closed BtuG lid, and there are no particle populations with open BtuG1 lids (Fig. 4b). We also collected data in the presence of 2 molar equivalents CNCbl, but 2D classes suggest that this led to destabilisation of the complex, similar to what we observed for BtuB2G2 (Fig. 4c).

**Fig. 4.**
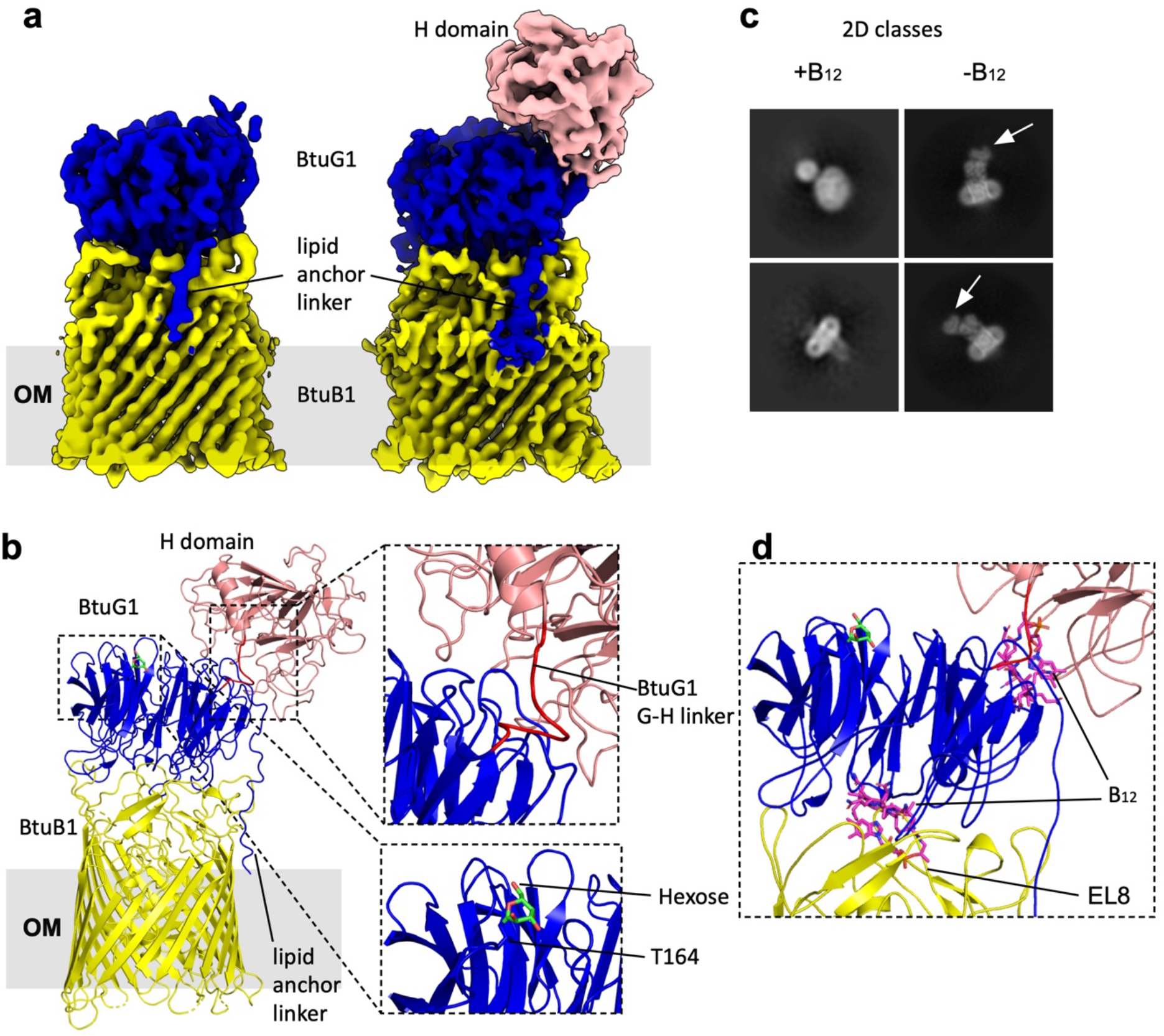
The cryo-EM structure of BtuB1G1 is closed. **a**, Cryo-EM maps used to build BtuB1 and BtuG1 G domain (left panel) and BtuG1 H domain (right panel). BtuB1 in yellow, BtuG1 G domain in blue and BtuG1 H domain in salmon. **b**, Cartoon model showing BtuG1 and BtuG1, colours as in (a). The linker between the domains G and H is shown in red for spatial reference; note it has been modelled into very poor density (top close up view). The bottom close-up view shows an O-Glycosylation site ^26^ for BtuG1 at residue Thr164. **c**, Representative 2D classification images from samples with and without B_12_. White arrows denote the BtuG1 H domain. **d**, Close-up view of a cartoon model for BtuB1G1 showing the two B_12_ binding sites occupied. B_12_ has been placed by superimposing BtuH2-B_12_ (7BIZ)^25^ and BtuG2-B_12_ onto BtuB1G1. Note the steric clashes of the modelled B_12_ molecules with BtuB EL8 and with the BtuG1 G domain.

### Insights into BtuB2G2 B_12_ lid opening from molecular dynamics

To quantify the opening of the lid, we selected the Cα atoms of Tyr37 from the BtuG2 linker and of Pro468 from BtuB2 EL7, whose relative positions remain stable during the simulations, and of Asp236 from BtuG2 located on the opposite site of the hinge, *i*.*e*., in the widest opening region (Fig. 5). The spatial relation of the three Cα atoms (two static and one mobile) allows calculation of an aperture angle α. The angle is approximately 18° in the crystal structure (Fig. 5a). Two unbiased simulations with different initial atomic velocities at T=300 K gave mixed results. In one 2 µs-long simulation there is an opening of up to α=25°, but in the other one the aperture remains similar to that in the crystal structure (Fig. 5b and Supplementary Fig. 8a). In the simulation where opening occurs, BtuG2 moves away from BtuB2 in a lid-like motion. EL2 of BtuB2 and the BtuG2 linker act as a hinge, and the partial opening is on the opposite side (Fig. 5). To further explore opening, we ran a 5 µs-long simulation (Supplementary Fig 8b-c). This revealed fluctuations up to α=28°, but no permanent lid opening. To assess whether the large number of hydrogen bonds (∼40) and salt bridges between BtuB2 and BtuG2 were hindering lid opening, we also ran two simulations at 400 K. In these simulations, we observe lid openings of up to α=37°, suggesting that lid opening requires energy input and/or longer time scales (Fig. 5c and Supplementary Fig. 8d). A free energy simulation shows that the energy associated with the opening of the lid has a minimum for α = 18-22° but quickly increases to exceptionally high values of over 40 kcal/mol for α > 40° (Fig. 5d). These energies might be slightly overestimated due to shortcomings of classical MD simulations^27^, but it is clear that opening angles of 40° and higher are highly unlikely at ambient temperatures on microsecond timescales in our *in silico* setup. It should be noted that, for the sake of simplicity, we used an OM composed of PL in our simulations. The presence of LPS in the OM could conceivably lower the stability of the closed state of the transporter, *e*.*g*. via interactions with BtuB ELs.

**Fig. 5.**
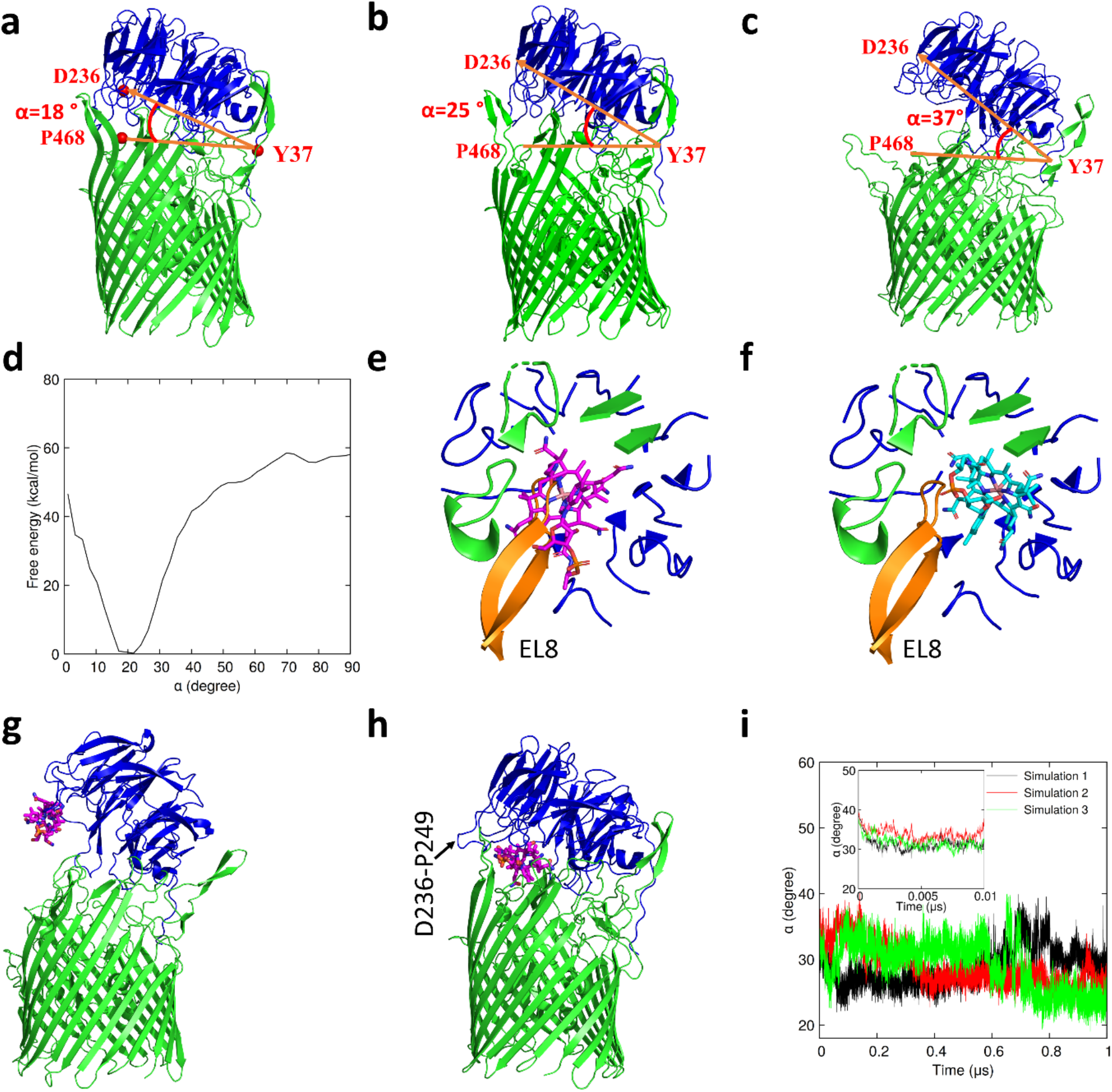
BtuG2 lid opening and BtuB2G2 CNCbl acquisition investigated by molecular dynamics. **a**, Definition of the opening angle in the crystal structure of BtuB2G2. See text for details. **b**, Unbiased simulations at 300 K reach an opening angle of 25°. **c**, Opening angle of 37° observed at 400 K. The time dependence of the angle is shown in Supplementary Fig. 7a,b and d. **d**, Free energy profile of the lid opening as a function of the angle α. **e**, Superimposition of the crystal structures of BtuG2-CNCbl and the BtuB2G2 complex revealed that the EL8 loop (shown in orange) of BtuB2 occupies the CNCbl binding cavity of BtuG2 in BtuB2G2 and would block access of CNCbl. **f**, The docked CNCbl is oriented in the binding cavity of BtuG2 without a steric clash with the EL8 loop. **g-h**, Structural changes of the BtuB2G2 complex and movement of the ligand during a 1 µs-long simulation of the CNCbl acquisition process. The starting and final conformations are shown in g and h, respectively. **i**, The angle α for three different simulations for CNCbl acquisition by BtuB2G2.

An important question is how substrate molecules enter the BtuB2G2 complex. Although spontaneous binding of CNCbl to BtuG2 was readily observed in the MD simulations (Fig. 2c), the stability of the closed BtuB2G2 structure would make it extremely unlikely to observe a spontaneous entering of CNCbl into an open BtuB2G2 complex. Thus, we set out to simulate the reverse “unbinding” process in simulations by applying an external force. To this end, we docked CNCbl into the BtuG2 binding pocket within the BtuB2G2 complex. It was not possible to simply place the CNCbl at its position in the crystal structure of BtuG2-B_12_ due to steric clashes with EL8 from BtuB2 (Fig. 5e). Therefore, we docked a CNCbl molecule into the BtuG2 binding pocket within BtuB2G2 to avoid those clashes (Fig. 5f and Supplementary Fig. 9a). We then pulled the CNCbl molecule away from the OM along the membrane normal, *i*.*e*. along the BtuB2 barrel axis, using steered MD (SMD) with a small constant force of 100 kJ/mol/nm^2^ (Supplementary Fig. 9a-d). The COM of the BtuB2 barrel was held fixed during these simulations. In this manner, the CNCbl molecule caused the BtuG2 lid to open and moved out of the BtuB2G2 complex. A snapshot of the simulation just before the CNCbl molecule loses contact with the BtuB2G2 complex is shown in Fig. 5g. In this simulation, the lid opens to an angle of about α=40° which is similar to the opening during unbiased simulations at 400 K (Fig. 5c). This model, with a substrate molecule close to the mouth of the partially open complex, served as input for the subsequent simulations of B_12_ acquisition by a partially open BtuB2G2 complex.

Three 1 µs-long unbiased MD simulations were performed with CNCbl at the mouth of the complex oriented as obtained from the SMD simulation. Two things happen spontaneously during these simulations: the CNCbl molecule rapidly moves into a position close to the BtuG2 binding pocket and the lid starts closing. Due to the different initial velocities in these simulations, the details of substrate movement and lid closing vary, but they happen in all three replicates. The position of the substrate reached after 1 µs is slightly different from that in the crystal structure of BtuG2-CNCbl due to the altered conformation of loop β5A (Supplementary Fig. 9e,f). Simulation 3 was particularly interesting; here, the BtuG lid closed very slowly up to 600 ns and after that, a quick CNCbl acquisition was observed together with a further closing of the lid to α ∼23° (Fig. 5i, green curve; Supplementary movie 2). While no major EL8 rearrangement was observed, a slight reorientation of this loop occurred in all three simulations to accommodate CNCbl in the binding cavity. The β5A loop moves very frequently during the simulations and adopted a final orientation similar to the one in the BtuB2G2 crystal structure. In an additional simulation (data not shown) using a randomly selected position and orientation of CNCbl close to the outer surface of BtuG2, ligand acquisition was not observed. It is likely that a proper orientation of the molecule has to be achieved before/during the movement toward the BtuG2 binding pocket within the partially open state, which might take a relatively long time. In any case, an analysis of the electrostatics of the (partial) open BtuB2G2 complex suggests that BtuG2 will still attract CNCbl to its binding site (Supplementary Fig. 10). Finally, to account for the possibility that the complex opens more than the 40 degrees achieved via the approaches described above, we also generated a wide-open state (α = ∼60°) via SMD. We then replaced BtuG2 with the BtuG2-CNCbl crystal structure, followed by a 1.5 μs unbiased simulation. In this case, however, the complex did not close due to the fact that the β5A loop remained bound to CNCbl (final α ∼ 35°; Supplementary Fig. 11a-c).

### CNCbl transfer from BtuG2 to BtuB2

To generate the allosteric signal that exposes the TonB box of BtuB2 for interaction with periplasmic TonB, the substrate needs to be transferred from BtuG2 to the binding site in BtuB2. Therefore, we next explored how CNCbl is transferred to BtuB2 via a 3.5 µs-long unbiased MD simulation (Supplementary Fig. 12). As starting structure, we used the CNCbl-BtuB2G2 docked model used for the SMD simulations. We also docked CNCbl into BtuB2 at a position very similar to that for *E. coli* BtuB (Supplementary Fig. 7) and used this as the target position ^28^. As a measure for the movement of CNCbl, the centre of mass (COM) distance between CNCbl in the BtuB2 binding pocket and the initial position of CNCbl in BtuG2 was used. The initial COM distance of CNCbl is 16 Å from the BtuB2 binding site. After 1.5 µs, the distance had decreased to 10 Å. During this time, CNCbl underwent a dissociation-association process from the BtuG2 cavity, adopting a tilted orientation stabilised by electrostatic, CH-π, and π-π interactions with BtuG2 and BtuB2. It should be noted that the interaction of BtuG2 with BtuB2 causes a conformational change in BtuG2, *i*.*e*. in the complex the β5A loop shifts away from where the CNCbl ligand would be. As shown earlier in the simulation of BtuG2-CNCbl, the β5A loop interacts with the ligand. In the BtuB2G2 complex, however, this loop cannot contact the ligand due to the presence of BtuB2 (Fig. 5h). Thus, the CNCbl molecule is likely to be less strongly bound to the BtuG2 binding site within the complex and therefore can spontaneously move towards the BtuB2 binding site. At the end of the simulation, the ligand has moved 8 Å closer to the BtuB2 binding site (Supplementary Fig. 12). The very stable Asn54-Thr64 loop of BtuG2 inhibits further translocation in the unbiased MD simulation, with CNCbl getting trapped close to the BtuB2 binding site due to strong interactions with, among others, Phe58 and Gln59 of BtuG2 and Asp377 and Tyr321 of BtuB2.

To obtain a statistically more meaningful picture of the complete translocation pathway from the BtuG2 to the BtuB2 binding sites, free energy simulations were performed using the Metadynamics approach, independent of the previous unbiased simulations. The results show that the CNCbl molecule can move toward the BtuB2 active site in a tilted manner and reorients itself within the BtuB2G2 cavity during this translocation process (Fig. 6a,b). The calculated free energy profile reveals that the BtuG2 and BtuB2 binding sites are separated by about 10 kcal/mol in free energy. Between these two binding sites, the CNCbl molecule moves more or less in a barrierless process even though in individual unbiased simulations, like the one described above, the molecule can get stuck at energy minima for longer times. In the biased free energy calculations, CNCbl is able to reorient and reach the BtuB2 binding site, which is about 10 kcal/mol energetically more favourable than that of BtuG2. As proposed above, the CNCbl movement away from BtuG2 is likely made possible by the lack of plier-like interactions between CNCbl and the β5A and Asn54-Thr64 loops in the BtuB2G2 complex.

**Fig. 6.**
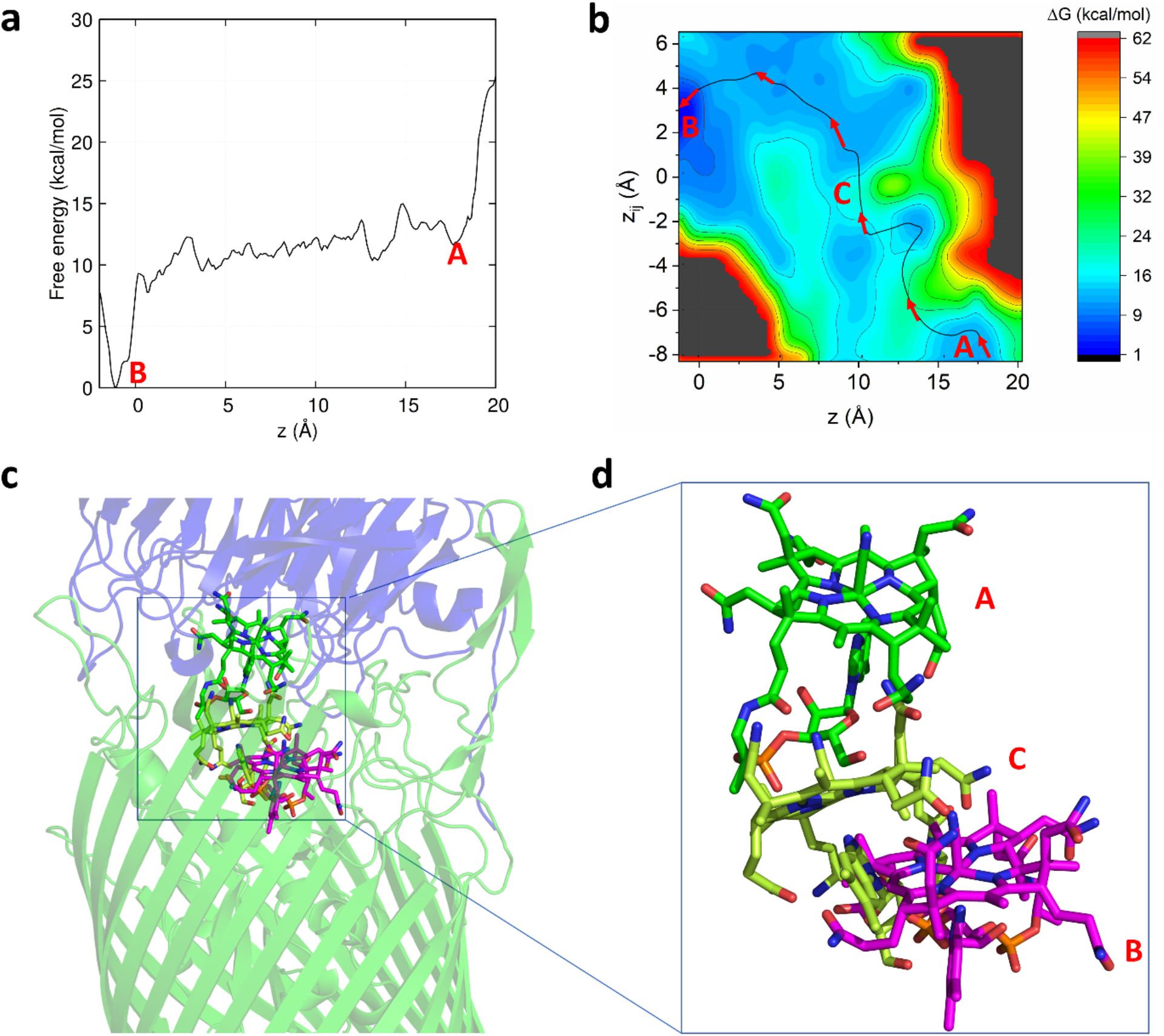
Free energy profile of the CNCbl translocation process. **a**, 1D free energy profile from the BtuG2 binding site (denoted by A) to the BtuB binding site (denoted by B). **b**, 2D free energy surface with the second collective variable *z*_*ij*_ indicating the orientation of the molecule (see Methods). The relatively flat energy landscape indicates that the CNCbl molecule can easily rotate when moving between the two binding cavities. **c-d**, Overview of the CNCbl translocation process (close-up view in d). CNCbl at the BtuG2 binding site (A) is in green and CNCbl in the BtuB2 binding site (B) is in magenta. An intermediate conformation (C) is shown in lime green.

An intriguing characteristic of many gut *Bacteroides spp*. is the presence of multiple B_12_ acquisition loci. This raises the important question as to whether these systems are redundant or different, complementing each other in order to ensure optimal B_12_ capture from the environment. An earlier study reported very different abilities of single BtuB mutants to compete with wild type *B. theta* when B_12_ is limiting. Strains encoding only BtuB1 or BtuB3 had large competitive defects, whereas a strain with only BtuB2 competed efficiently with wild type. In addition, cobalamins with different lower ligands selected for different single-BtuB strains in competition assays^18^. While these results could suggest different substrate specificities for the BtuB transporters, there is no information on OM levels of the various transporters (and indeed for any component of the three B_12_ loci), and how those levels could be affected by different cobalamins. For example, the apparent efficiency of BtuB2 could easily be explained by a much higher level within the OM relative to BtuB1 or BtuB3. To start addressing this important question, we performed quantitative OM proteomics on log- and stationary phase wild type *B. theta* grown in B_12_-limiting conditions (0.4 nM CNCbl) or B_12_-replete conditions (40 nM CNCbl). As shown in Fig. 7a, components of all three loci are present in the OM at low CNCbl concentration. Ranking the proteins on their relative label-free absolute protein quantification (iBAQ)^29^ intensities, an approximation of absolute protein abundance in a sample, under limiting B_12_, BtuB2G2 is the most highly abundant, followed by BtuB1G1. BtuB3G3 may be present at lower numbers in the OM. These data suggest that the higher fitness of BtuB2 might be due to higher amounts present in the OM^18^. The recently described BtuH proteins, which are surface-exposed and also bind B_12_ with high affinity^25^ appear to be more abundant than BtuB and BtuG. Interestingly, the uncharacterised Bt1491 protein of locus 1 is likely the most abundant protein by a considerably margin across all three loci under B_12_-limiting and B_12_-replete conditions, making it a priority to assess the potential role of this protein in B_12_ acquisition. Its locus 2 paralog Bt1957, on the other hand, has a very low iBAQ ranking (Fig. 7a). Both Bt1491 and Bt1957 have signal sequences and a cysteine around position 20-30 followed closely by several acidic residues, consistent with surface exposure^30^. Finally, a comparison of the iBAQ ranking in B_12_-limiting conditions vs. B_12_-replete conditions suggests that all components of the three loci are much more abundant in the OM under limiting B_12_ (Fig. 7b), as expected from transcriptomics data and the fact that the loci are under control of B_12_-responsive riboswitches^18,31^. During B_12_-replete conditions (40 nM CNCbl), expression of all Btu proteins is very much reduced, and most of the proteins were undetectable by mass spectrometry in OM samples (Fig. 7c). For future work, it will be interesting to assess the effect of different upper and lower ligands on OM protein levels, and determine absolute copy numbers of the various proteins per cell.

**Fig. 7.**
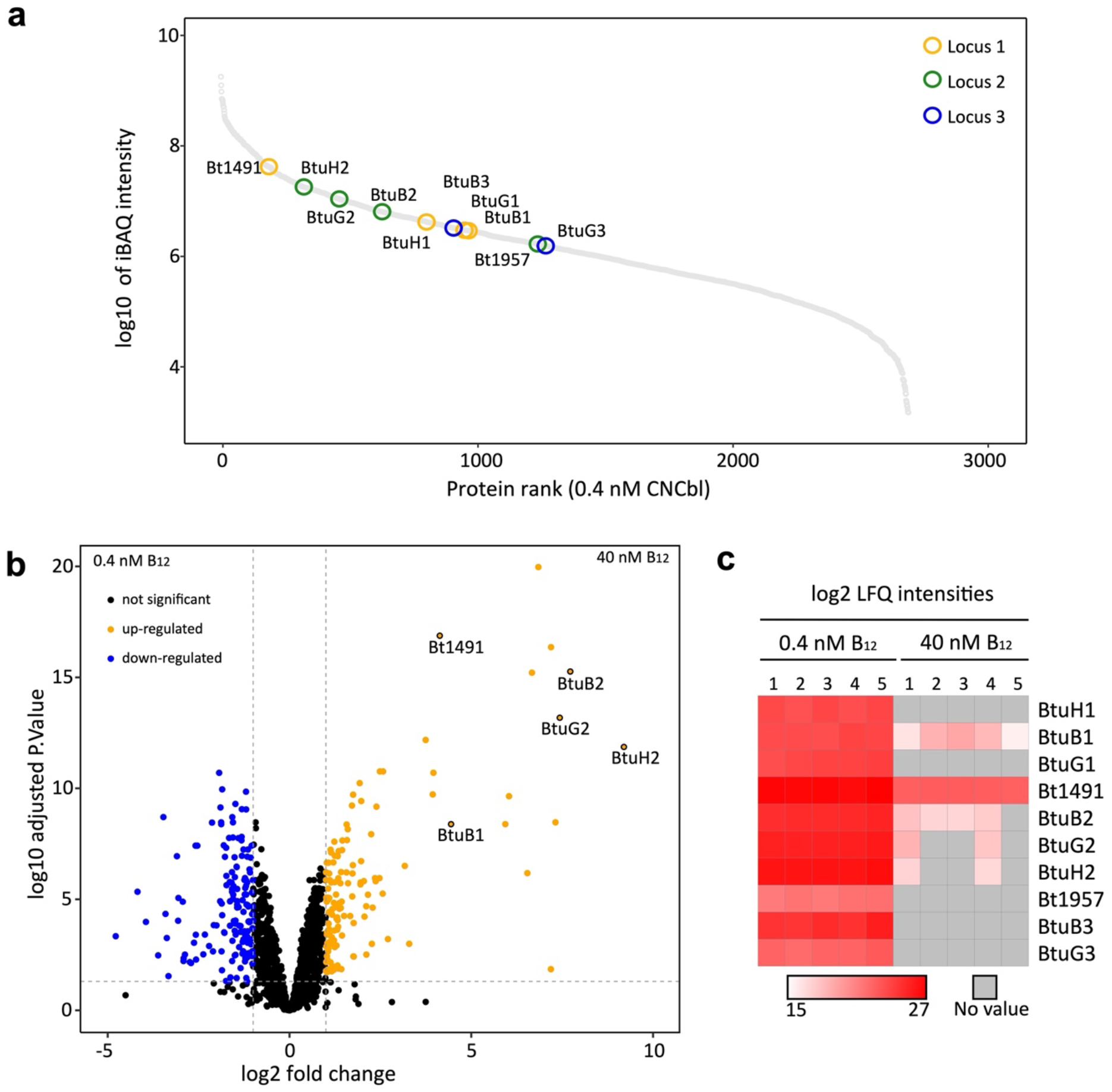
Proteomic analysis of *B*.*theta* OM under B_12_-restrictive (0.4 nM CNCbl) vs. B_12_-replete conditions (40nM CNCbl.) **a**, Relative abundance of OM proteins calculated using iBAQ intensities for cells grown in B_12_-restrictive conditions. **b**, Volcano plot showing the magnitude of change in the expression of OM proteins between B_12_-restrictive vs. B_12_-permissive conditions. Note that several proteins from the three loci are not detected under at least one condition. **c**, Heatmap showing the LFQ intensities in both conditions, with each square representing an independent sample. Five independent cultures were grown in minimal media supplemented with 0.4 or 40 nM CNCbl for 18 hours in anaerobic conditions. Grey indicates the protein has not been detected in this condition.

## DISCUSSION

How do the transporters and additional surface-exposed B_12_-binding lipoproteins such as BtuH, and possibly others, work together to take up corrinoids? The BtuBG transporters are reminiscent of the widespread SusCD systems that mediate OM uptake of complex glycans ^21,32^. Both complexes likely work in a similar fashion, consist of a TBDT with a mobile cap (SusD or BtuG), and use a “pedal-bin” mechanism for substrate loading and import^21,32^. However, in contrast to SusCD systems for which open and closed states are observed via cryo-EM, our data only show closed BtuBG complexes. In addition, loading purified BtuBG with B_12_ is challenging, again contrasting with SusCD systems that readily and stably bind substrate. Given the location of the BtuG B_12_ binding sites, it is clear that the BtuBG pedal bins will have to open in order for BtuG to capture ligand. Apparently, the energy landscapes for lid opening are very different for BtuBG and SusCD systems, with deep minima for BtuBG closed states. One possibility is that the stably closed BtuBG complexes result from OM extraction and purification in detergent. It is conceivable that the LPS within the native OM environment, being highly negatively charged^33^, could destabilise the closed state to favour opening. Alternatively, BtuG lid opening could be a relatively rare event, given that the B_12_ requirements of the cell are likely to be modest, contrasting with SusCD systems which mediate uptake of carbon sources. Another possibility is that the additional B_12_-binding surface lipoproteins, such as BtuH, are involved in loading the BtuBG complexes, *e*.*g*. via generation of the open transporter.

How are additional B_12_-binding surface lipoproteins like BtuH organised relative to the core BtuBG transporter? Again, a comparison with SusCDs provides some insights. SusCs are atypical TBDTs in that they are ∼50% larger than regular TBDTs such as BtuB (∼120 vs ∼75 kDa). In a very recent study, it was shown that one function of the increased SusC size is to provide binding interfaces for the additional surface lipoproteins, in this case glycoside hydrolases and surface glycan binding proteins. Thus, multiple OM components of a polysaccharide utilisation locus (PUL) are organised within one large complex on the cell surface, termed utilisome ^34^. BtuB proteins do not have the extra interfaces, and there is no room for stable association of other lipoproteins with the BtuBG core, which agrees with our pull downs assays in which only BtuB and BtuG co-purify, even in mild detergents. Thus, any interaction between BtuBG and other B_12_-binding lipoproteins is likely to be transient. This then leads to a picture that contrast sharply with utilisomes, with B_12_-binding lipoproteins in the OM that associate transiently with their cognate BtuBG transporter. This also raises the question whether those lipoproteins could function in an “inter-locus” manner, *e*.*g*. could BtuH2 assist vitamin B_12_ uptake by BtuB3G3?

Why would such differences in lipoprotein association exist? We speculate that this is due to the nature of the substrates and cellular requirements. Glycans are high molecular weight carbon sources that need to be processed into smaller fragments before being transported, while B_12_ requires no processing. To ensure efficient capture of the processed glycans, it would make sense for the processing machinery (*i*.*e*. the additional lipoproteins) to be closely associated with the transporter, which would ensure relatively high turnover numbers. By contrast, the main purpose of B_12_-binding lipoproteins would be to ensure capture of a relatively rare and valuable ligand without the need to be closely associated with the transporter to ensure high turnover. Indeed, the picomolar binding affinity of BtuH2^25^ results in part from a very low k_off_ (∼8.6 × 10^−5^ s^-1^), corresponding to an average lifetime of the B_12_-BtuH2 complex of several hours. Thus, on bacterial time scales, B_12_ is bound virtually irreversibly and will likely be released only after interaction of BtuH2 with BtuB2G2. The low k_off_ might also compensate for the low mobility of OMPs and LPS that emerges from studies in *E. coli*^35,36^. The fundamental difference in the nature of the substrate (polymers for SusCDs, a relatively small molecule for BtuBG) most likely explains why SusCD systems are dimers ^21^ and BtuBG monomers. Within the dimeric levan utilisome, the glycan binding sites of the surface glycan binding proteins are ∼100 Å apart, and such dual “anchor points” might allow for more efficient processing of a polymer ^34^.

Related to the point discussed for BtuH2 above, free BtuG2 has an even higher affinity for CNCbl (∼10^−13^ M), and the average lifetime of the B_12_-BtuG2 complex is about an hour^19^. This is obviously also too long to be meaningful *in vivo*. Our MD simulations show that upon closure of the complex, one of the extracellular BtuB2 loops prevents the β5A loop of BtuG2 to clamp B_12_, and this would likely result in a lower affinity for the ligand and faster k_off_. Thus, we propose that steric clashes in the BtuG lid that result from pedal bin closure cause conformational changes in the lid that lower the affinity for the ligand, followed by transfer to BtuB (Fig. 8). For SusCD systems, such conformational changes have not been observed and indeed seem not necessary since SusD lids bind their substrates with much lower affinities (μM-mM). In conclusion, lipoprotein-capped TonB-dependent transporters seem to be exclusive for Bacteroidetes and most likely offer a competitive advantage in the gut. Previous data for SusCD complexes, added to our current characterisation of the BtuBG systems of *B. theta*, suggest that the majority of *Bacteroides* TBDTs function in a lipoprotein-assisted manner.

**Fig. 8.**
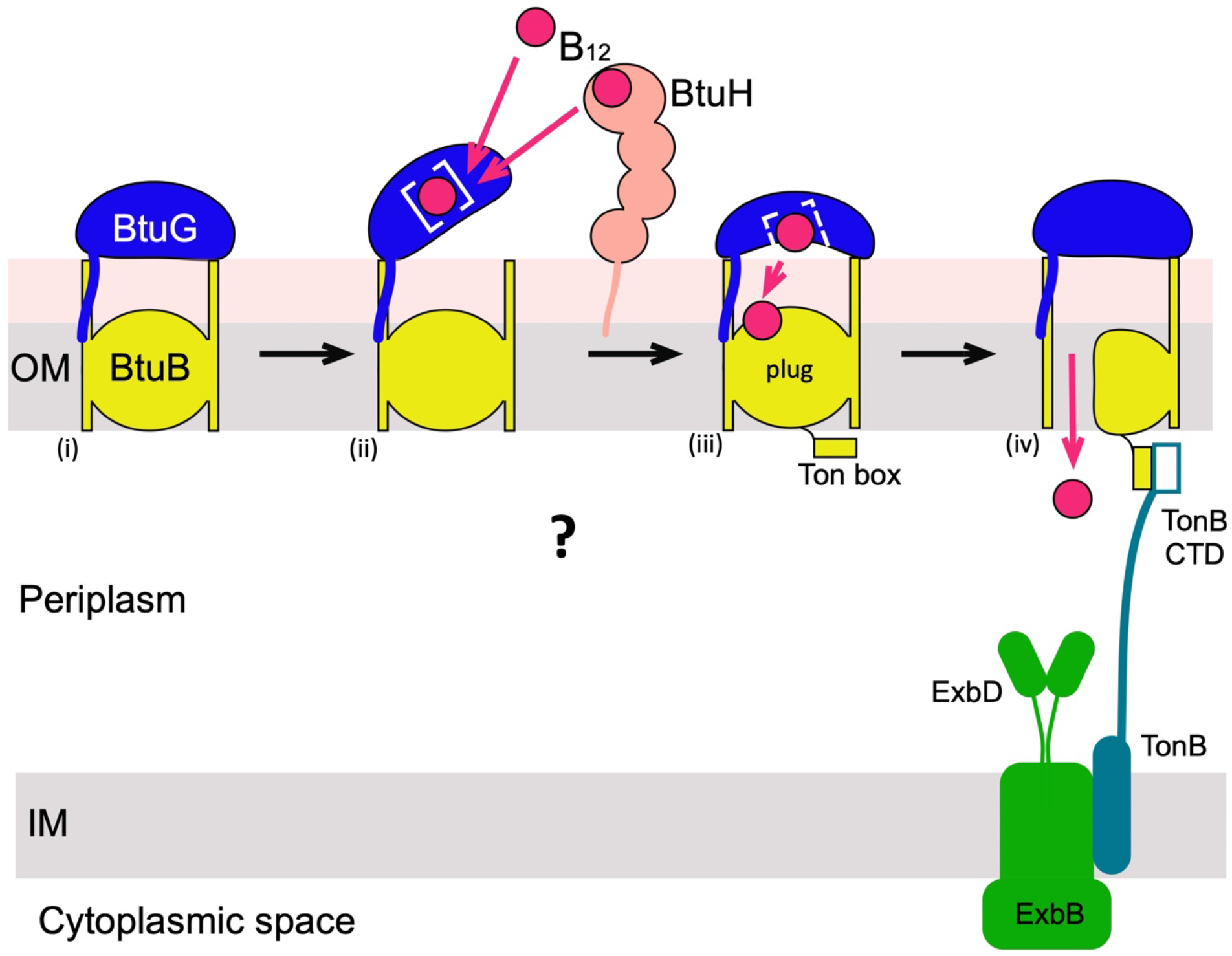
Schematic model for lipoprotein-mediated B_12_ acquisition by *B. theta*. **The** first stage of acquisition is the opening of the BtuBG complex, which might be promoted by LPS (pink) or accessory proteins such as BtuH. (ii) B_12_ would be transferred from BtuH to BtuG or bound from the extracellular space by BtuG directly. (iii) Upon closing, BtuG undergoes conformational changes to avoid clashes with BtuB, causing the release of B_12_ and its transfer to BtuB. Binding of B_12_ by BtuB generates allosteric changes in the plug that leads to TonB box exposure in the periplasmic space. During the final stage (iv), the C-terminal domain (CTD) of TonB binds to the TonB box and causes unfolding of the plug due to mechanical force generated by the TonB-ExbB-ExbD complex in the IM. The channel that is formed allows diffusion of the substrate into the periplasmic space ^37^.

## METHODS

### Cloning, expression and protein purification

The coding region for the mature peptide of *bt1954 (btuG2)* was cloned into pET28 using NcoI and XhoI. The protein was expressed in BL21(DE3) 2.5h at 37°C with the addition of 1mM IPTG. Cells were collected by centrifugation, resuspended in TBS (10mM Tris, 300 mM NaCl; pH8) lysed at a pressure of 20-23Kpsi using a cell disrupter, and purified by nickel affinity chromatography. Concentration of the protein was measured using the BCA assay, and when needed, adenosylcobalamin, cyanocobalamin or cobinamide was added to a ratio 1:2 protein:corrinoid. The sample was incubated at 4°C for 30 minutes and further purified using a HiLoad 26/600 Superdex 200 (GE healthcare).

### Over-expression of BtuB2

To a strain with loci 1 and 3 deleted^18^ a 6 x his-tag was added to the C terminus of genomic *bt1953* (*btuB2*) using pExchange-*tdk*. The expression of locus 2 was driven by P1E6 and not the B_12_-dependent wild type riboswitch. The mutated strain was grown for 20 hours in minimal media with 0.5% fructose under anaerobic conditions and collected by centrifugation at 11,000g for 30 minutes. Pellets were resuspended in TBS and lysed at a pressure of 23 Kpsi using a cell disrupter (Constant Systems 0.75kW). The membrane fraction was harvested by centrifugation (45 minutes at 42,000 r.p.m. using a 45 Ti Beckman rotor). The inner membrane from 4 litres was solubilized in 0.5% sodium N-lauroyl sarcosine in 20 mM HEPES pH 7.5 and the sample was centrifugated for 30 minutes at 42,000 r.p.m in a 45 Ti Beckman rotor. The pellet was solubilized in TBS plus 1.5% n-dodecyl-N,N-dimethylamine-N-oxide (LDAO anatrace) 1h at 4°C and centrifuged again. The supernatant was loaded onto a nickel column (TBS 0.2% LDAO 25mM imidazole) and the eluted sample (250 mM imidazole) further purified by size exclusion chromatography (HiLoad 26/600 Superdex 200 GE healthcare) using 10 mM HEPES pH 7.5, 100 mM NaCl and 0.05% LDAO (anagrade). The eluted samples from 10 purifications (∼2.5 mg total) were pooled together and buffered exchanged into 10 mM HEPES pH 7.5, 100 mM NaCl and 0.4% tetraethylene glycol monooctyl ether (C_8_E_4_).

### Over-expression of BtuB1

We introduced a C-terminal hexa-his tag into *BT1489* (*btuB1*) in a *B. theta* mutant strain lacking loci 2 and 3^18^. We then replaced the wild type promoter (B_12_-dependent riboswitch) with the constitutive promoter P1E6 to obtain high levels of expression. Growth conditions were similar to BtuB2 but the minimal media was supplemented with 0.25 mM L-methionine (as the strain lacks both inner membrane B_12_ ABC transporters). Five 4l batches of cells were purified in the same way as for BtuB2 with a minor modification, the last size exclusion chromatography was run using 10 mM HEPES pH 7.5, 100 mM NaCl and 0.05% DDM. The yield obtained was ∼50 micrograms/l.

### Structure determination

Sitting drop vapour diffusion crystallization trials were set up with a Mosquito Crystallization robot (TTP Labtech) using the commercial screens MemGold1 and MemGold2 (Molecular Dimensions) for BtuB2-BtuG2 (in the presence of 1.5-fold molar excess of CNCbl) and JCGS+, Structure (Molecular Dimensions) and Index (Hampton research) for BtuG2. For BtuG2-CNCbl and BtuG2-AdoCbl we obtained crystals in the condition E4 from JCSG+ (0.2M Lithium sulphate, 0.1M Tris pH 8.5, 1.26 M ammonium sulphate). These were cryoprotected with saturated ammonium sulphate. For BtuG2-Cbi crystal were obtained in C9 from Index (1.1M Na Malonate pH7, 0.1M HEPES pH7 and 0.5% Jeffamine ED2001 pH7) and cryoprotected with 20% PEG400. For the complex BtuB2-BtuG2 the initial hit C2 from MG2 (0.08M magnesium acetate tetrahydrate, 0.1M sodium citrate pH 6.0, 14% PEG 5000 MME) was further optimised with variations of 2% of the original PEG concentration. The crystals were cryoprotected using 20% PEG400 and flash frozen in liquid nitrogen. Diffraction data were collected at 100K at Diamond Light Source (supplementary Table1). Dataset for the Co-SAD experiment was integrated using XDS^38^, pointless^39^ was used for space group determination and then scaled and merged with Aimless^39,40^. To find the heavy atom substructure, phenix AutoSol^41,42^ was used. Clear electron density was visible for BtuG2 and for a molecule of B_12_. An initial model was built using AutoBuild^43^ in phenix and manual building in coot. The high-resolution data set for BtuG2-CNCbl was integrated with Dials^44^, while BtuG2-AdoCbl and BtuG2-Cbi were integrated with XDS^38^. These three datasets were scaled and merged using Aimless^39,40^. The initial model produced from the Co-SAD experiment was used to solve the phase problem by molecular replacement with Phaser^45^. All the models were improved by rounds of manual building using coot^46^ and refinement using Refmac 5.8^47^ in CCP4 cloud^48^ for BtuG2-AdoCbl and BtuG2-CNCbl and Phenix refine^49^ for the Co-SAD experiment and BtuG2-Cbi. Models for CNCbl (CNC code in coot) AdoCbl (B1Z) and Cbi (CBY) were placed in coot. For the high-resolution model anisotropic refinement with automated local NCS (non-crystallographic symmetry) restrains were used while isotropic refinement with TLS and automated local NCS restrains were selected for BtuG2-AdoB_12_.

Diffraction data for the BtuB2-BtuG2 complex was integrated with XDS^38^, the space group was determined with pointless, then scaled and merged with Aimless^39,40^. The phase problem was solved with two components search in Phaser^45^ using the models BtuG2 (HR) and 2GUF (the BtuB structure from Escherichia coli Sculptor-modified within Phenix). The model was built with iterative cycles of AutoBuild^43^ and manual building in coot^46^. The model was refined in Phenix using secondary structure and BtuG2 model restrains, NCS and TLS. MolProbity^50^ was used to validate protein geometry and PyMol was used for the visualization of the protein structures. To calculate the evolutionary conservation of the amino acid positions, the phylogenetic relations of 150 homologous sequenced were analysed and colour coded according to their conservation value using ConSurf server^23^. The multiple sequence alignment was built using MAFFT, collecting the homologues from UNIREF90. The homolog search algorithm used was HMMER (HMMER-value: 0.0001) with 1 iteration and a maximal and minimal percentage of identity of 95% and 35% respectively. The chains used for the analysis were C and D from the BtuB2-BtuG2 model.

### Cryo-EM structure determination

Purified BtuB1G1 complex (∼2.5 mg/ml) without CNCbl was applied to glow-discharged Quantifoil 0.6/1 300 mesh holey carbon grids, blotted and plunge-frozen in liquid ethane using a Vitrobot Mark IV device (Thermo Fisher Scientific). Grids were imaged on a FEI Titan Krios microscope (Thermo Fisher Scientific) operating at 300 kV (Supplementary Table 2). Movies were recorded using EPU (Thermo Fisher Scientific) on a Falcon 4 direct electron detector in electron event representation (EER) mode at 130,000 magnification, corresponding to a pixel size of 0.91 Å. The microscope was equipped with a Selectris X energy filter with slit width set to 5 eV. 1,924 movies were recorded in total. Data were processed in cryoSPARC v3.3.2^51^ (Supplementary Table 2). Following patch motion correction and patch CTF correction, ∼2,000 particles were manually picked and used to make templates for template-based picking. 1,110,283 particles were extracted in 300 × 300 pixel boxes and subjected to two rounds of 2D classification. Good classes were selected and used to make an ab initio model using a stochastic gradient descent approach with 2 classes. The class showing secondary structure features was subjected to non-uniform refinement^52^. Several rounds of heterogeneous refinement, non-uniform refinement and 2D classification were used to discard the remaining bad particles, resulting in the final particle stack with 50,547 particles. The particles were re-extracted in 384 × 384 pixel boxes and subjected to a final round of non-uniform refinement with defocus and global CTF refinement enabled, resulting in a 3.22 Å reconstruction. An *ab initio* model was built into the cryo-EM map using Buccaneer (part of the CCPEM package^53^) followed by iterative manual building in Coot^46^ and real space refinement in phenix^54^. The initial atomic model for BtuB1 and the G domain of BtuG1 was used to sharpen the density using LocScale^55^ to improve interpretability and aid with model building. To build the H domain of BtuG1, the final reconstruction from non-uniform refinement was sharpened with a B-factor of –50 Å^2^. The BtuB1G1 model built into the LocScale sharpened map was refined into the B-factor-sharpened map together with the H domain of BtuG1. There is strong density extending from the side chain of T164 located on the β-propeller of BtuG1, which likely corresponds to the presence of a glycan moiety. The precise composition of the O-glycan is unknown, therefore a single β-D-galactopyranosyl modification was modelled. The refinement statistics of the final model can be found in Supplementary Table 2.

Purified BtuB1G1 complex with 2 equivalents CNCbl was applied to glow discharged Quantifoil 0.6/1 300 mesh holey carbon grids, blotted and plunge frozen using a Vitrobot Mark IV device. The grids were imaged on a FEI Titan Krios microscope operating at 300 kV. Movies were recorded on a Falcon 3 direct electron detector in counting mode at 75,000 magnification, corresponding to a pixel size of 1.065 Å. 626 movies were recorded in total. Data were initially processed as above, but following 2D classification no 3D reconstruction with secondary structure features could be obtained from *ab initio* reconstructions or non-uniform refinement.

### MD simulation methods

All BtuB2G2 systems for the MD simulations in the presence and absence of cyanocobalamin were built using the CHARMM-GUI Membrane Builder^56^. Using the PROPKA webserver, the protonation states of the titratable amino acids were verified to be in their standard protonation states. The BtuB2G2 protein was placed into a 1-palmitoyl 2-oleoyl phosphatidyl-ethanolamine (POPE) bilayer and solvated with TIP3P water molecules on both sides of the membrane maintaining a water thickness of about 25 Å. Since demineralized water is a typical artefact of MD simulations and as salt can greatly influence protein stability and conformations, we studied the proteins under physiological conditions of 0.15 M KCl by adding K^+^ or Cl^-^ ions to achieve the desired concentration and to neutralize the system. The total system composed of the protein, membrane, solvent, ions, and ligand was placed into a rectangular box of 80 × 80 × 150 Å^3^. For the MD simulation of CNCbl-bound-BtuB2G2, the CNCbl was docked to the binding pocket of BtuG2 and BtuB2 using the Autodock program. A cyanocobalamin ligand parameter file for the CHARMM program was available in the literature and converted to the GROMACS format^57^. For simulations in the absence of BtuB2, the surface-exposed lipoprotein BtuG2 was simulated in an aqueous environment after neutralizing the 18*e* negative charge with K^+^ ions and in the presence of additional 0.15 M KCl. The system composed of BtuG2, water, ions, and ligand was placed into a rectangular box of size 72 × 88 × 77 Å^3^. All simulations were conducted using the GROMACS molecular dynamics software, version 5.1.14, and employing the CHARMM36-m forcefield^58,59^.

Energy minimization for the model systems was performed using the steepest descent method followed by a two-step constant volume (NVT) equilibration of 5ns each with varying restrains on the simulated system at 300 K. The NVT simulations were carried out using a Berendsen thermostat with a 1-picosecond temperature coupling constant. Furthermore, the systems were relaxed in a four-step constant pressure (NPT) equilibration of 50 ns by removing the restraints on the protein, the membrane, and the ligands in a stepwise manner. The NPT simulations were carried out using the semi-isotropic coupling method to a Berendsen barostat at one bar with a coupling constant of 2 ps. For non-bonded interactions, the Verlet cut-off scheme for the Coulomb and Lennard-Jones interactions was employed with a cut-off of 10 Å. Moreover, the Particle Mesh Ewald scheme was used for evaluating the long-range electrostatic interactions, and all bonds in the proteins were constrained using the Linear Constraint Solver (LINCS) algorithm^60,61^. After the six-step equilibration, the final simulations were continued according to the protocols defined for unbiased and steered molecular dynamics (SMD) simulations. The NPT production runs were performed using a Parrinello-Rahman^62^ barostat along with a semi-isotropic pressure coupling and the Nose-Hover thermostat^63^ (unbiased, SMD, and metadynamics MD simulations). The steered MD simulations were performed by pulling the center of mass of CNCbl and BtuG2 protein along the channel axis with constant velocity of 1 Å/ns using a spring constant of 100 kJ/mol/ns^2^. To understand the system stability and structural changes during the permeation of cyanocobalamin through the BtuBG protein, we analysed the root mean square deviations from the initial structure (RMSD) and the radius of gyration (R_g_) of the proteins (data not shown). Further angle analyses were performed to examine the extent of lid opening of the BtuBG protein complex via a hinge loop. The short-range electrostatic interaction between the CNCbl and BtuG was calculated using the ‘*gmx energy*’ tool.

### Free energy calculations

To determine the free energies of biomolecular processes, the metadynamics (Mtd) technique was employed that progressively builds up a history-dependent biasing potential along predefined collective variables (CVs)^64^. The sum of these Gaussian potentials is given by^65^

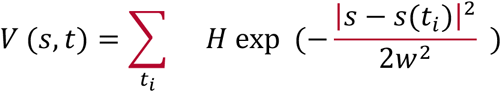

where *H* denotes the height of the Gaussian hills, *w* their width, and s the CV. In a MtD simulation, the bias or “hills” are dynamically placed on top of the underlying potential energy landscape and discourage the system from re-visiting the same points in configurational space. This extra bias enforces, e.g., an unbinding process, within a reasonable computational time. In the well-tempered version of metadynamics, which is used in the study, one rescales the Gaussian height with the bias accumulated over time at a fictitious higher temperature, T+ΔT^66^. Despite its inherent nonequilibrium characteristics, one can extract information close to the true equilibrium state of the system by suitably tuning the parameter ΔT. The free energy profile usually termed potential of mean force (PMF) can be estimated as

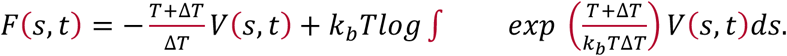

The center of mass between the CNCbl and the BtuG2 binding cavities name z was considered as the major CV for the binding free energy calculation between the CNCbl and BtuG2. The corresponding dissociation constant was calculated based on the free energy difference ΔG was estimated using *k* = *e*^−Δ*G*/*RT*^ at 300 K. To construct the 2D free energy surface, a second CV was defined as the projection *z*_*ij*_ of the distance *r*_*ij*_ between two porphyrin carbon onto the z axis (**Figure S3d**). The orientation of the substrate molecule can be determined as *φ* = *cos*^−1^(*r*_*ij*_/*z*_*ij*_), i.e., the value of *z*_*ij*_ determines the orientation of the molecule. To examine the free energy associated with the BtuG2 lid opening process, multiple-walker WTMTD simulations were carried out. The 1D free energy profile was estimated as a function of the CV angle, which can depict the aperture between ButG2 and BtuB2, between amino acids group Leu467-Gly469 of BtuB2, Asp36-Gly38 from the linker of BtuG2, and Thr235-Gly237 from the junction of a flexible and stable loop of BtuG2. Furthermore, the free energy associated with the CNCbl translocation from the BtuG2 binding pocket to the BtuB2 binding pocket was calculated using the COM distance between the CNCbl and binding cavity of BtuB2 as biasing CV, z, and the distance between the z-component of the two porphyrin carbon atoms as reweighting CV (**Figure S3d**).

### Sample preparation and mass spectrometry for proteomics

Five independent cultures were grown in minimal media with 0.4 nM or 40 nM CNCbl and grown for 18 hours. Cells were collected by centrifugation at 11,000 ×g for 30 min. The pellets were resuspended in TBS and sonicated. The membrane fraction was harvested by centrifugation (45 minutes at 45,000 r.p.m. using a 45 Ti Beckman rotor). The fractions were washed two times with water (resuspended in water followed by 45 minutes at 42,000 r.p.m. using a 70 Ti Beckman rotor). Finally, they were stored at -80°C until needed.

Membrane fractions were subjected to denaturation and tryptic digest using a suspension trapping-S-trap (Protifi) protocol. Briefly, this included resuspension in 5 % SDS, 50 mM Tris pH 7.4, denaturation with 5 mM tris(2-carboxyethyl)phosphine (TCEP) at 60 °C for 15 min, alkylation with 10 mM N-ethylmaleimide (NEM) at room temperature for 15 min, and acidification to a final concentration of 2.7 % phosphoric acid. Samples were then diluted eightfold with 90 % MeOH 10 % TEAB (pH 7.2) and added to the S-trap micro columns. The manufacturer-provided protocol was then followed, with a total of five washes in 90 % MeOH 10 % TEAB (pH 7.2), and trypsin added at a ratio of 1:10 enzyme:protein (2 μg : 20 μg) and digestion performed for 2 h at 47 °C. Peptides were dried and stored at -80 °C, and immediately before mass spectrometry were resuspended in 0.1 % formic acid. LC-MS/MS was performed using an Ultimate 3000 RSLCnano System (ThermoFisher Scientific) in line with an Orbitrap Fusion Lumos Tribrid mass spectrometer (ThermoFisher Scientific). Peptides (1 μg) were injected onto a PepMap100 C18 LC trap column (300 μm ID x 5 mm, 5μm, 100 Å), and separated with an EASY-Spray nanoLC C18 column (75 μm ID x 50 cm, 2 μm, 100 Å) at a flow rate of 250 nl/min, column temperature 45 °C. Solvent A was 0.1 % (v/v) formic acid in HPLC water, and solvent B was 0.1 % (v/v) formic acid and 80 % (v/v) acetonitrile in HPLC water. LC-MS/MS runs were preceded by a 2-min equilibration with solvent A and solvent B at 98 % and 2 %, which was maintained for 5 min following sample injection, then the gradient increased solvent B to 35 % over the next 120 min. Solvent B was increased to 90 % in 30 sec for 4 min, then decreased to 2 % in 30 seconds to allow further equilibration for 10 min. Data were acquired by the Orbitrap Fusion Lumos in positive ion mode, with data-dependent acquisition. MS1 was performed at 120,000 resolution, in the scan range 400-1600 m/z, charge states 2-7, AGC target of 200,000, with a maximum injection of 50 ms with repeat count 1. Peptides were fragmented using HCD (30 % collision energy). The ion trap was selected as the detector type, set to rapid scan rate, mass range and scan range mode set to normal and auto, AGC target set to standard, maximum injection time set to dynamic. The entire duty cycle lasted 3 seconds, during which time “TopSpeed” analysis was performed.

### Proteome data analysis

Raw files from mass spectrometry were matched against the *B. thetaiotaomicron* reference proteome (UP000001414, downloaded from UniProt on 01/12/2021), and proteins quantified using MaxQuant V 2.0.3.0 ^67^. NEM alkylation of cysteine was set as a fixed modification, oxidation of methionine and acetylation of protein N-termini were set as variable modifications. Digestion was trypsin/p specific, LFQ and matching between runs was selected. All other parameters were left as defaults. The output from MaxQuant was processed in Perseus^68^, where contaminants and decoys were removed and iBAQ was used to rank protein expression within each condition. Processing was also performed using the Limma package ^69^ in the R programming environment, where LFQ intensity values were used to quantify relative protein abundance between different conditions. Proteins were filtered to remove contaminants and decoys, and those identified by <2 unique peptides. Changes in protein abundance between conditions were considered significant where there was a difference of at least twofold, and when Student’s T-test p-value =<0.05 after Benjamini-Hochberg correction for multiple comparisons. We used the R^70^ package Ggplot2^71^ to generate proteomic analysis graphs. Morpheus (https://software.broadinstitute.org/morpheus) was used to generate the heatmap.

## Data availability

The mass spectrometry proteomics data have been deposited to the ProteomeXchange Consortium^72^ via the PRIDE partner repository^73^ with the data set identifier: PXD0XXXX. Reviewer account details: Username: XXXXX Password: XXXXX

## Acknowledgements

The research of B.v.d.B is supported by a Wellcome Trust Investigator award (214222/Z/18/Z), providing salary support for J.A.R and A.S. We would like to acknowledge the Diamond Light Source for crystallography beam line access (proposal mx-24948) and i04 beamline support, Dr Dan Maskell and the Astbury Biostructure Laboratory (Leeds) for collecting cryo-EM data, the Newcastle Structural Biology Laboratory for data processing infrastructure and Professor Andrew Goodman (Yale University) for providing *Bacteroides thetaiotaomicron* strains. K.J acknowledges the Alexander von Humboldt (AvH) foundation for an AvH postdoctoral research fellowship. K.J and U.K are grateful to the North German Supercomputing Alliance (Norddeutscher Verbund für Hoch-und Höchstleistungsrechnen – HLRN) for providing access to their high-performance computational facilities. K.J thanks Jigneshkumar Dahyabhai Prajapati and Vinaya Kumar Golla for helpful scientific discussions. A.F and M.T are funded by a Wellcome Investigator Award to M.T (215542/Z/19/Z).

## Author contributions

J.A-R and B.v.d.B expressed and purified proteins and determined X-ray crystal structures. A.S determined the cryo-EM structure. K.J. and U.K. carried out molecular dynamic simulations.

A.B maintained the Newcastle Structural Biology Laboratory. B.v.d.B and U.K supervised the structural and computational studies, respectively. J.A-R, K.J, U.K and B.v.d.B contributed to writing the manuscript. A.F performed and analysed the proteomic experiments. M.T supervised the proteomic studies.

