## Supplementary Data file for "BtuB TonB-dependent transporters and BtuG surface lipoproteins form stable complexes for vitamin B_12_ uptake in gut *Bacteroides*"

**Supplementary Table 1 | X-ray crystallographic data collection and refinement statistics**. Values in parentheses are for the highest resolution shell

| | BtuG2-CNCbl Co-SAD | BTuG2-CNCbl hi-res | BtuG2-AdoCbl | BtuG2-Cbi | BtuB2G2 |
|---|---|---|---|---|---|
| **Data collection** | | | | | |
| DLS beamline | i24 | i24 | i03 | i24 | i04 |
| Wavelength | 1.54987 | 0.96858 | 0.89843 | 0.99987 | 0.97950 |
| Space Group | I222 | $P2_1$ | $P2_1$ | $P4_12_12$ | C2 |
| Cell dimensions | | | | | |
| a,b,c (Å) | 59, 129, 141 | 80, 59, 101 | 80, 58, 101 | 79, 79, 155 | 280, 154, 107 |
| a,b,g (°) | 90, 90, 90 | 90, 103, 90 | 90, 104, 90 | 90, 90, 90 | 90, 106, 90 |
| Molecules in AU | 1 | 2 | 2 | 1 | 3:3 |
| Resolution range(Å) | 64.6-1.9 (2.0-1.9) | 59.1-1.7 (1.7-1.7) | 70.3-2.3 (2.4-2.3) | 43.30-1.33 (1.36-1.33) | 76.3-3.7 (3.8-3.7) |
| I/ sI | 12.2 (1.3) | 6 (1.4) | 5.1 (1.5) | 16.7 (0.9) | 4.2 (1) |
| Completeness (%) | 99.9 (99.9) | 99.4 (96.8) | 100 (100) | 99.9 (98.3) | 99.3 (99.8) |
| Redundancy | 32.5 (26.7) | 3.5 (3.6) | 6.8 (6.5) | 24.2 (17.9) | 3.8 (3.9) |
| Rpim (%) | 4 (53.1) | 7 (51) | 10 (44) | 2.1 (82) | 16 (85) |
| CC (1/2) | 1 (0.69) | 0.98 (0.56) | 0.98 (0.69) | 1 (0.3) | 0.9 (0.3) |
| Anomalous completeness | 99.8 (99.8) | - | - | - | - |
| Anomalous redundancy | 15.9 (12.9) | - | - | - | - |
| **Phasing** | | | | | |
| SOLVE FOM | 0.25 | - | - | - | - |
| Sites found [expected] | 1 [1] | - | - | - | - |
| **Refinement** | | | | | |
| Resolution (Å) | - | 59.1-1.72 | 70.2-2.3 | 43.3-1.33 | 67.02-3.72 |
| Rwork/Rfree (%) | - | 17.9/22.3 | 22.3/26.8 | 14.2/16.8 | 23.9/29.1 |
| Reflections | - | 97918 | 41442 | 113069 | 45634 |
| No. Atoms | - | | | | |
| Protein | - | 5489 | 5454 | 2817 | 23359 |
| Corrinoid | - | 186 | 218 | 68 | - |
| B-factors (Å2) | - | | | | |
| Protein | - | 20.3 | 36.3 | 23.4 | 123.8 |
| Corrinoid | - | 16.8 | 27.5 | 19.6 | - |
| Rmsd | - | | | | |
| Bond lengths (Å) | - | 0.007 | 0.012 | 0.012 | 0.004 |
| Bond Angles (°) | - | 1.7 | 1.8 | 1.2 | 1.07 |
| Molprobity clashscore | - | 2.4 | 4 | 1.1 | 13.5 |
| Ramachandran plot | - | | | | |
| Favoured (%) | - | 93.7 | 92.3 | 93.2 | 91.2 |
| Disallowed (%) | - | 0 | 0.9 | 0.3 | 0.49 |
| PDB code | - | 8BMX | 8BMY | 8BMZ | 8BM0 |

**Supplementary Table 2 | Cryo-EM data collection, image processing and refinement statistics for BtuB1G1**

| **Data collection** | |
|---|---|
| Electron microscope | FEI Titan Krios |
| Voltage (kV) | 300 |
| Spherical aberration (µm) | 2.7 |
| Camera | Falcon 4 (counting) |
| Energy filter | Selectris X (5 eV slit) |
| Magnification | 130,000 |
| Pixel size (Å) | 0.91 |
| Total dose ($e^-$/Å$^2$) | 35 |
| Dose rate ($e^-$/pixel/s) | 6.22 |
| Defocus minimum maximum (µm) | -0.9 to -2.4 |
| Number of EPU frames | 160 |
| Number of movies collected | 1,924 |
| **Image Processing** | |
| Initial number of particles | 1,110,283 |
| Final number of particles | 50,547 |
| Global resolution (FSC = 0.143) | 3.22 |
| Map sharpening | B-factor (–50 Å$^2$) |
| **Refinement** | |
| Model composition | |
| Non-hydrogen atoms | 9883 |
| Protein residues | 1250 |
| R.m.s. deviations | |
| Bonds lengths (Å) | 0.003 |
| Bond angles (˚) | 0.547 |
| Validation | |
| Molprobity score | 2.31 |
| Clash score | 14.83 |
| Rotamer outliers (%) | 0 |
| Ramachandran plot | |
| Favoured (%) | 86.51 |
| Outliers (%) | 0.08 |
| PDB | 8BLW |
| EMDB | EMD-16114 |

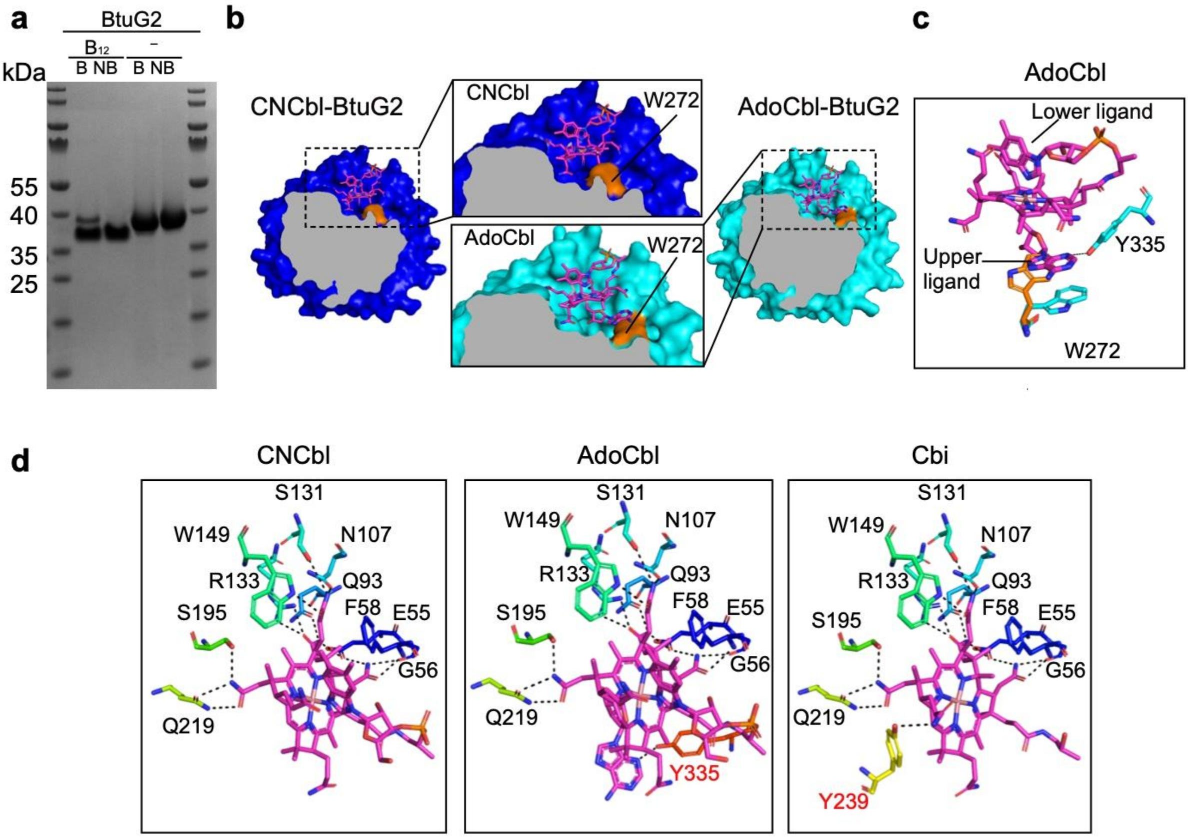


**Supplementary Figure 1 l BtuG2 binds different corrinoids**. **a**, Gel showing the different mobility of BtuG2 bound to CNCbl ($B_{12}$). B stands for boiled, NB for non-boiled. Notice the double band on the CNCbl-BtuG2 boiled sample; the higher band corresponds to apo-BtuG2 after $B_{12}$ removal due to the boiling. **b**, Slice of a surface representation for CNCbl-BtuG2 (blue) and AdoCbl-BtuG2. Close up views to show the cavity in which the upper ligand (-CN or -5'deoxydenosyl respectively) is located. W272 is depicted in orange; this residue is displaced in AdoCbl-BtuG2, enlarging the binding pocket. **c**, Residues implicated in the upper ligand stabilization of AdoCbl (coloured in cyan). The orange W272 corresponds to the position of the side chain in the CNCbl-BtuG2 structure. **d**, Panels displaying the residues involved in hydrogen bonding between BtuG2 and different corrinoids seen in the crystal structures. Common residues for the binding of the three corrinoids have black labels, those implicated in only one of the corrinoids red. Residues in rainbow colouring are ramped from blue at the N terminus to red at the C terminus. Hydrogen bonds are represented as black dashed lines.

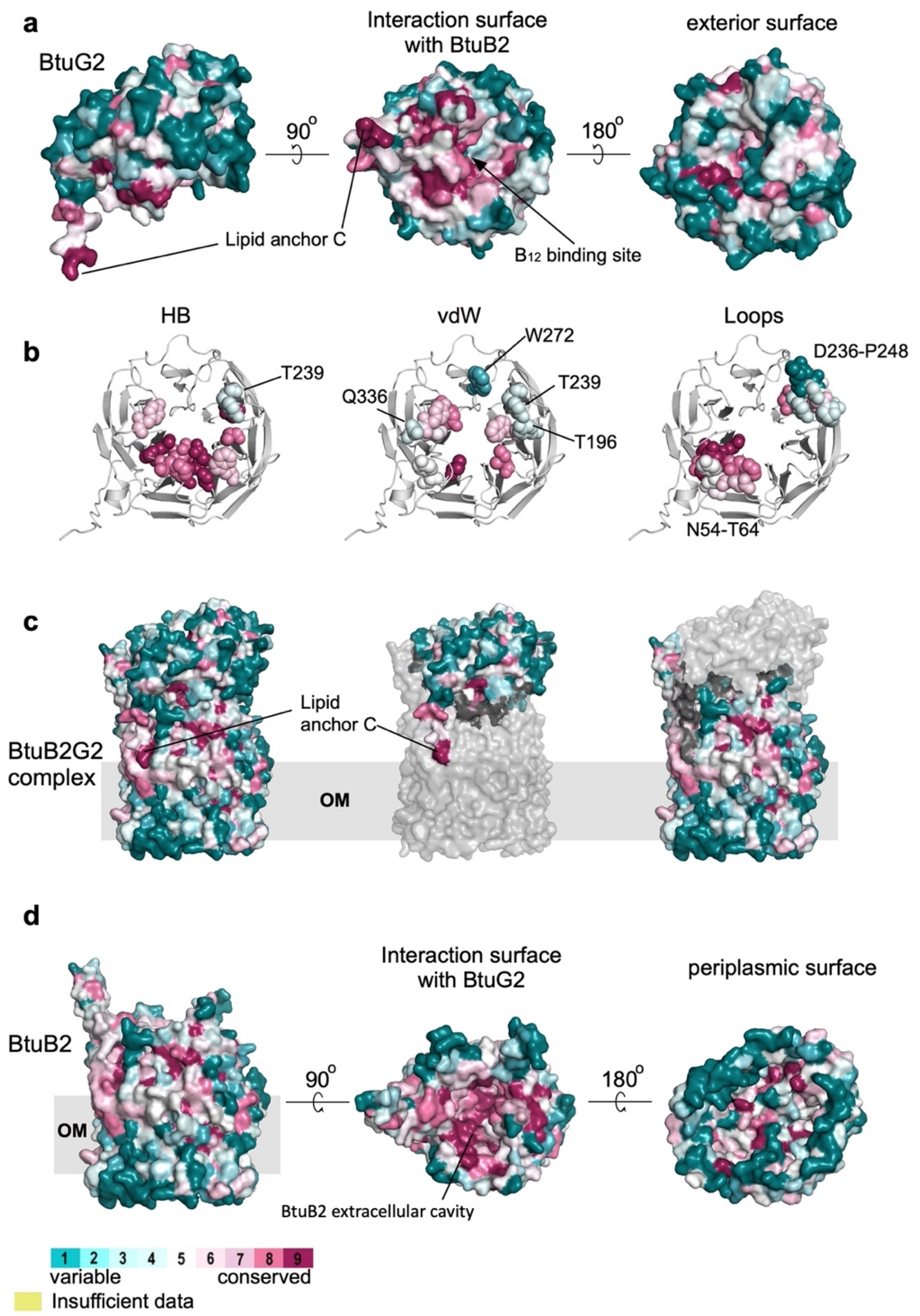


**Supplementary Figure 2 l ConSurf analyses of BtuG2 and BtuB2**. **a,** Several views of the surface representation of BtuG2 coloured with the conservation scores. **b,** Cartoon representation of BtuG2 showing the level of conservation of residues implicated in hydrogen Bonding (left panel), van der Waals forces (middle panel), and interacting loops derived from the structural and molecular dynamic simulations (right panel) implicated in the ligand binding. **c,** Surface representation of BtuB2-BtuG2 complex with the conservation score colouring. In the middle panel BtuB2 is in grey and in the right panel BtuG2 is grey. **d,** Similar to (a), but for BtuB2.

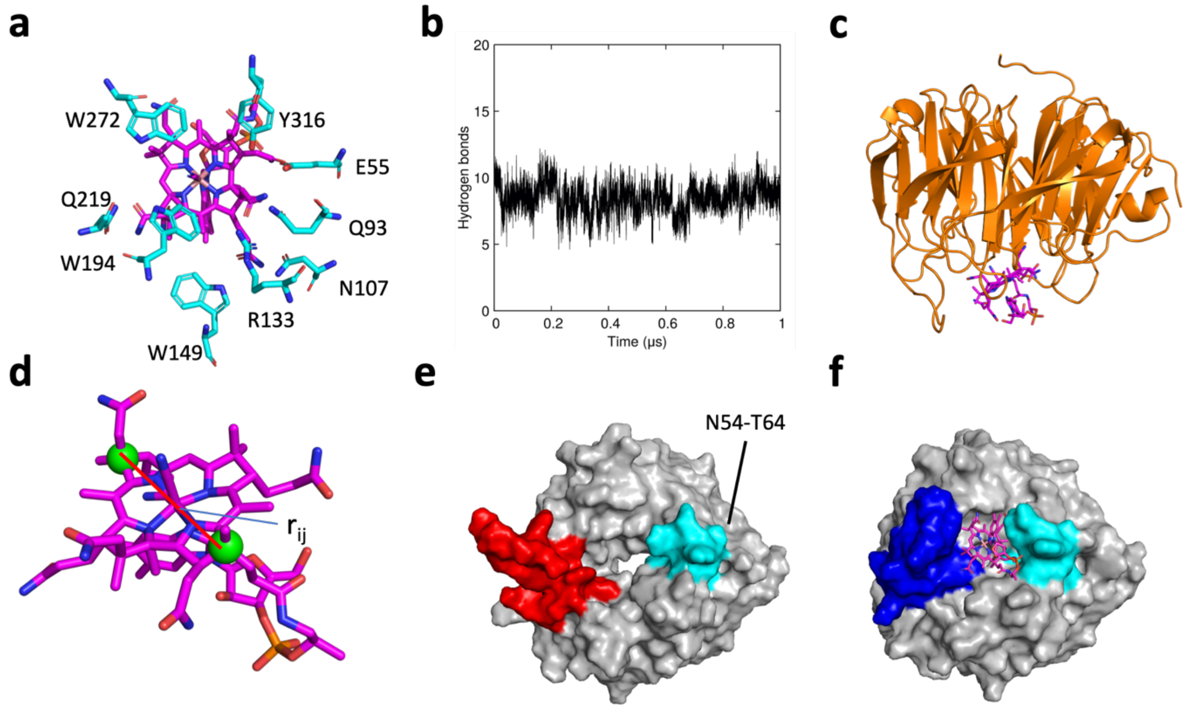


**Supplementary Figure 3 l Analysis of CNCbl binding by BtuG2 using MD simulations. a,** BtuG2 amino acid residues within 3Å of the COM of the binding site of CNCbl based on the crystal structure. **b,** Hydrogen bond interactions between the residues of the COM and CNCbl during the simulation. **c,** Representative snapshot of the CNCbl-BtuG2 at the end of the 1 μs-long MD simulation performed using the BtuG2-CNCbl crystal structure. **d,** Projection onto the z axis of the distance $r_{ij}$ between the carbon atoms depicted as pink spheres is defined as collective variable (CV) $z_{ij}$. **e-f**, Surface representation showing loops β5A (D236-P249) and N54-T64 from the relaxed state (left panel; β5A red) and crystal structure (right panel; β5A blue). Loop N54-T64 is coloured cyan.

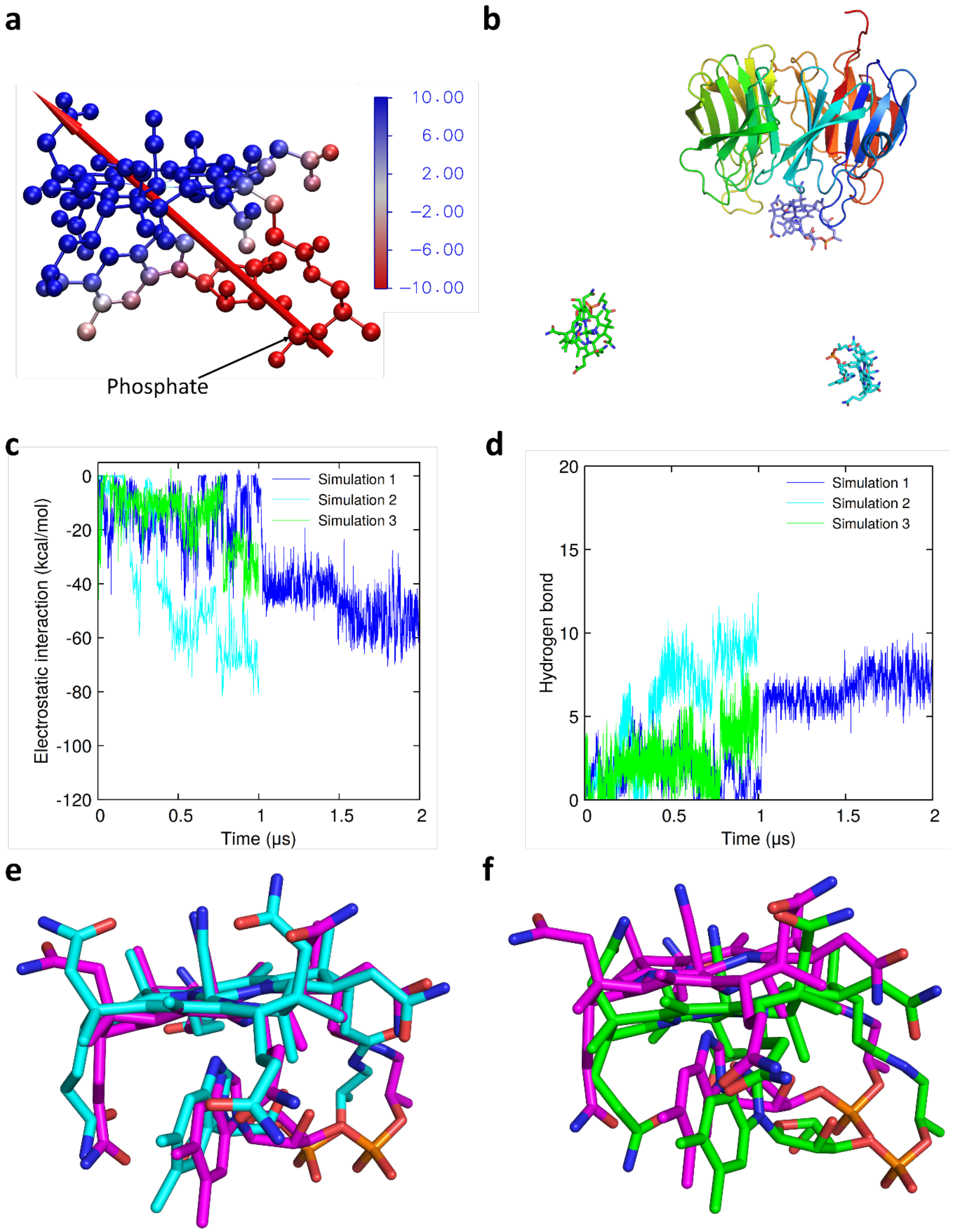


**Supplementary Figure 4 l Dipole of CNCbl and CNCbl acquisition by BtuG2 from different starting locations. a,** To show the dipole character of CNBl, the electrostatic potential projected onto the molecular structure is shown. The corrin ring is partially positive, whereas the nucleotide loop, including the phosphate group, is negatively charged. **b,** Representation of the initial positions of CNCbl (8.5, 37.1, and 34.9 Å away from the binding pocket for simulation 1 (dark blue), 2 (light blue) and 3 (green), respectively). **c,** Short-range electrostatic interaction energies between the ligand and protein throughout the simulations. **d,** Hydrogen bond interactions between residues of the active site and CNCbl. For comparison, there are 10 hydrogen bonds present in the crystal structure. **e**, The final position of CNCbl obtained from the unbiased simulation 2 (cyan carbons) is compared to the crystal structure position (magenta). A representative overlay of the final ligand position for simulation 1 is shown in Fig 2d. **f**, Same as **e** but for simulation 3 (green carbons).

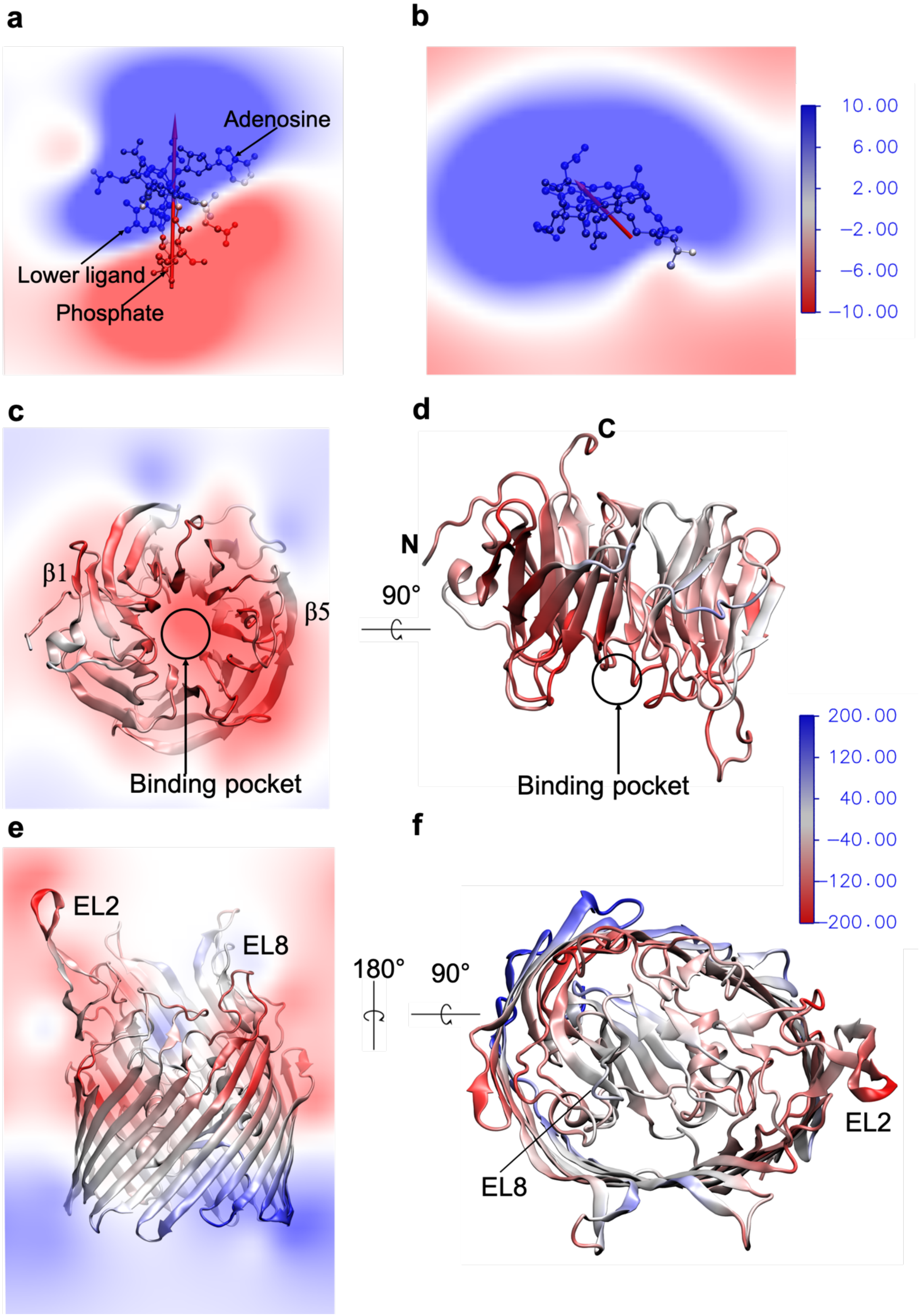


**Supplementary Figure 5 l Electrostatic potential maps. a,** Electrostatic potential maps of the neutral AdoCbl molecule with a large dipole moment. The corrin moiety of AdoCbl is partially positive whereas the lower ligand is partially negatively charged. **b,** Electrostatic potential maps of the positively charged Cbi. The colour scale is shown from -10, negative surface (red), to 10, positive surface (blue) in units of $k_B T/e$ = 26 mV at 300 K. **c-d,** The electrostatic potential map of BtuG2 is predominantly negative because of the net charge of -18*e*, explaining why BtuG2 attracts the positively charged or zwitterionic CNCbl/AdoCbl/Cbi. The BtuG2 in this figure has been obtained by removing the CNCbl from the CNCbl-bound-BtuG2 crystal structure. The colour scale is shown from -200, negative surface (red), to 200, positive surface (blue). **e-f,** The electrostatic potential map of BtuB2. The colour scale is same as for **(c-d)**.

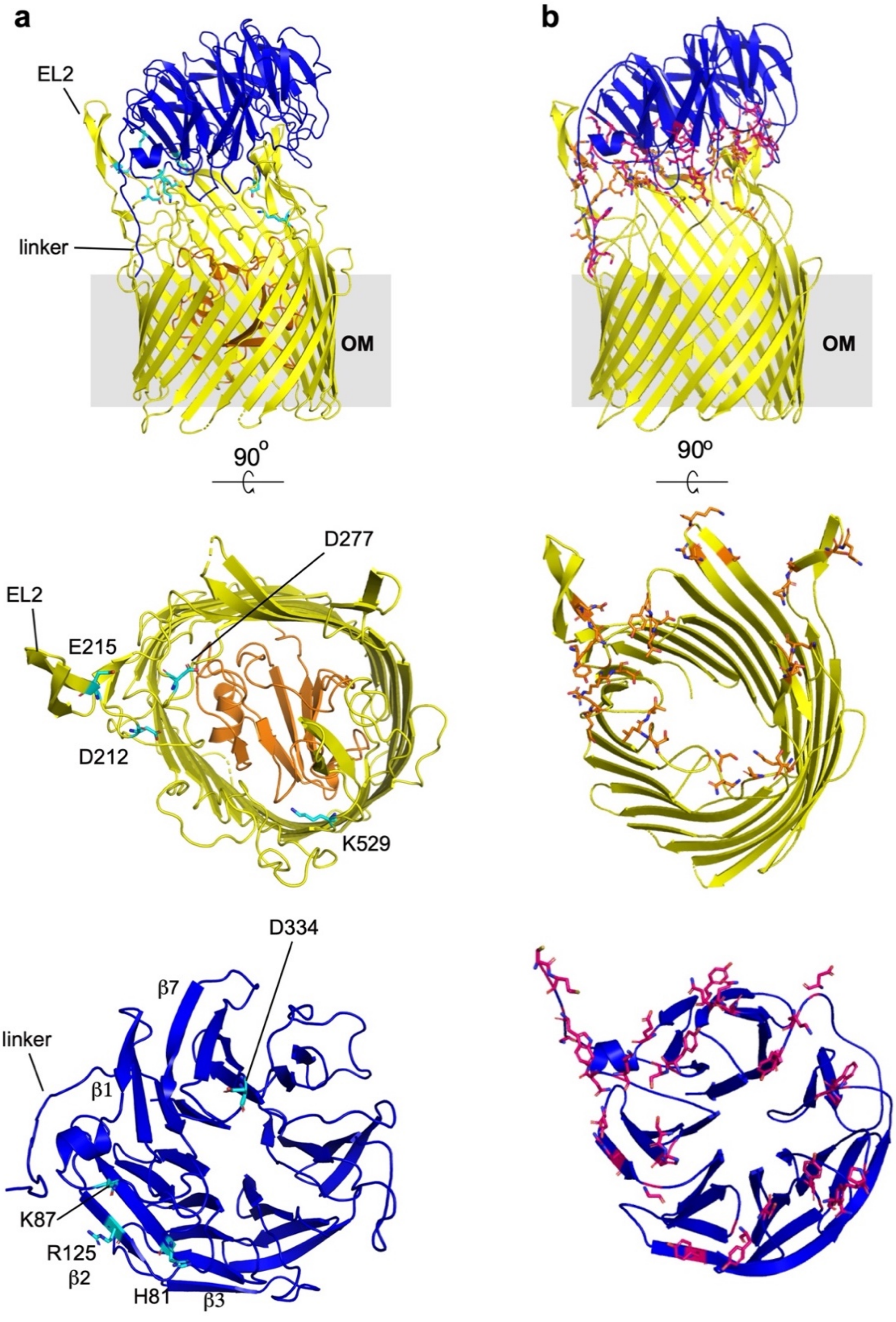


**Supplementary Figure 6 l Electrostatic interactions between BtuG2 and BtuB2. a**, Cartoon representations showing residues implicated in salt bridges in cyan. BtuB2 is in yellow (plug in orange) and BtuG2 in blue. **b**, Residues implicated in hydrogen bonding in red for BtuG2 and orange for BtuB2. The bottom panels show the BtuG2 surface viewed from the direction of BtuB2. For clarity, the BtuB2 plug has been removed and loops are smoothened in b.

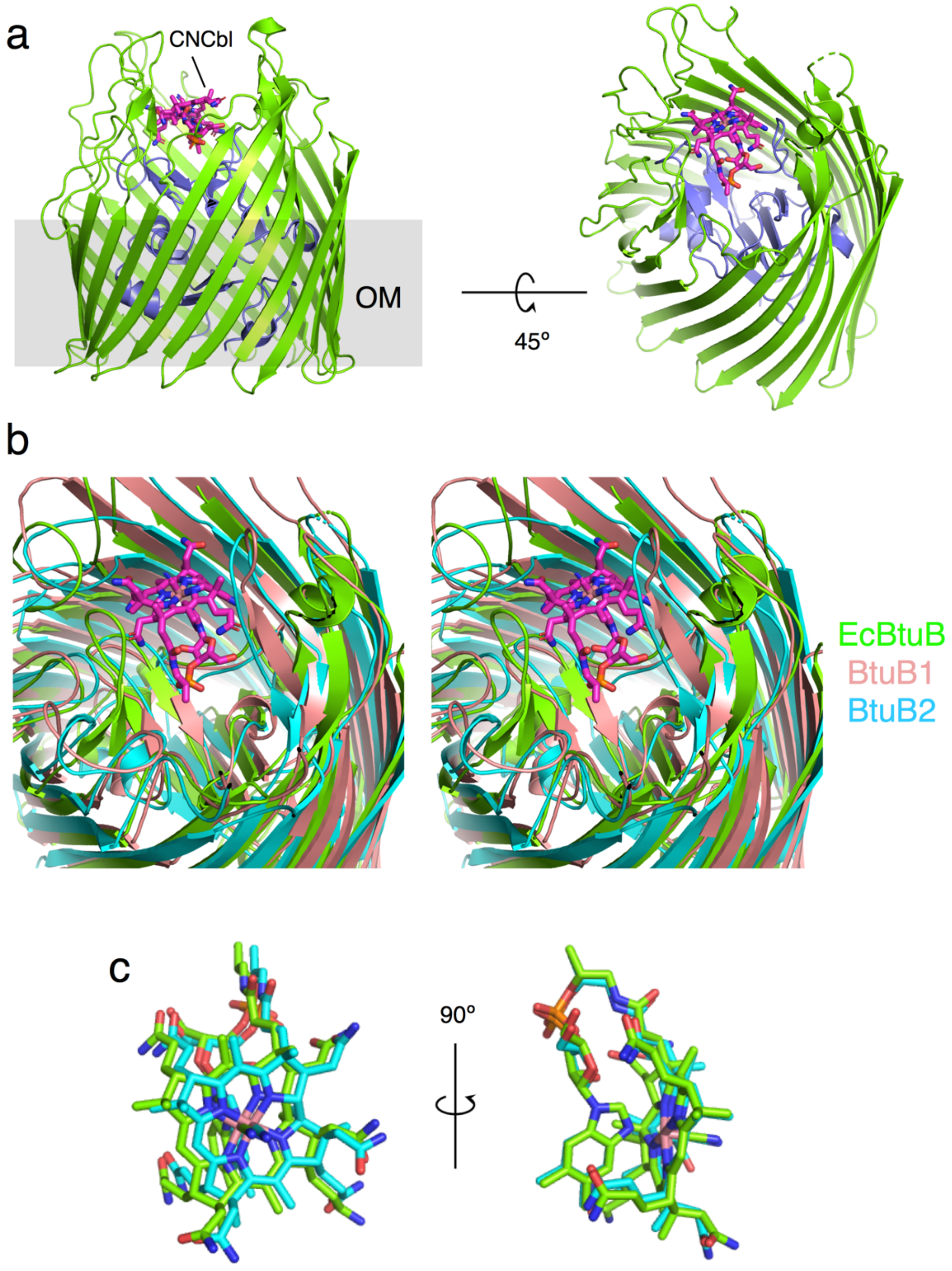


**Supplementary Figure 7 | Comparison of B12 binding sites in different BtuBs. a**, Cartoon model of EcBtuB viewed from the OM plane (left) and from the outside of the cell, with bound CNCbl shown as a stick model. The EcBtuB plug is slate blue. **b**, Stereo cartoon superpositions of EcBtuB and *B. theta* BtuB1 and BtuB2, showing the similarity of the $B_{12}$ binding sites. Loops have been smoothened for clarity. **c**, Comparison of bound CNCbl in EcBtuB with docked CNCbl in BtuB2. The views were generated via superposition of the BtuB proteins.

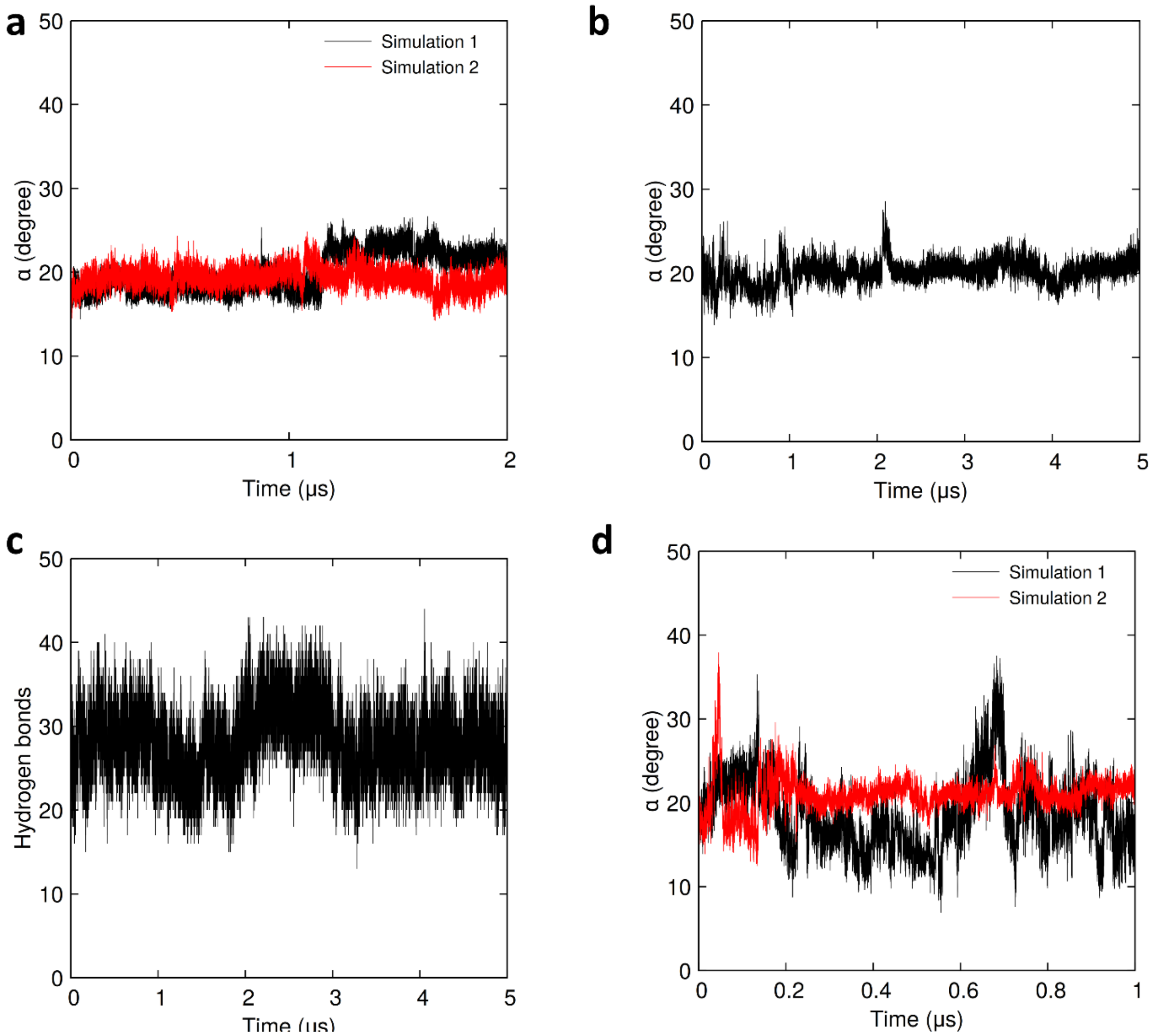


**Supplementary Figure 8 | Conformational changes of the BtuB2G2 complex during unbiased simulations. a**, Representation of the aperture angle, α, for 2 μs simulations at 300 K. The starting point is the closed state seen in the crystal structure (angle α of 18°). **b**, Same as in (a) but for the 5 μs unbiased simulation. **c**, Average number of hydrogen bonding interactions between BtuB2 and BtuG2. **d**, Analysis of the aperture angle for two simulations run at 400K; note the short-duration maximum aperture of around 37° observed in both simulations.

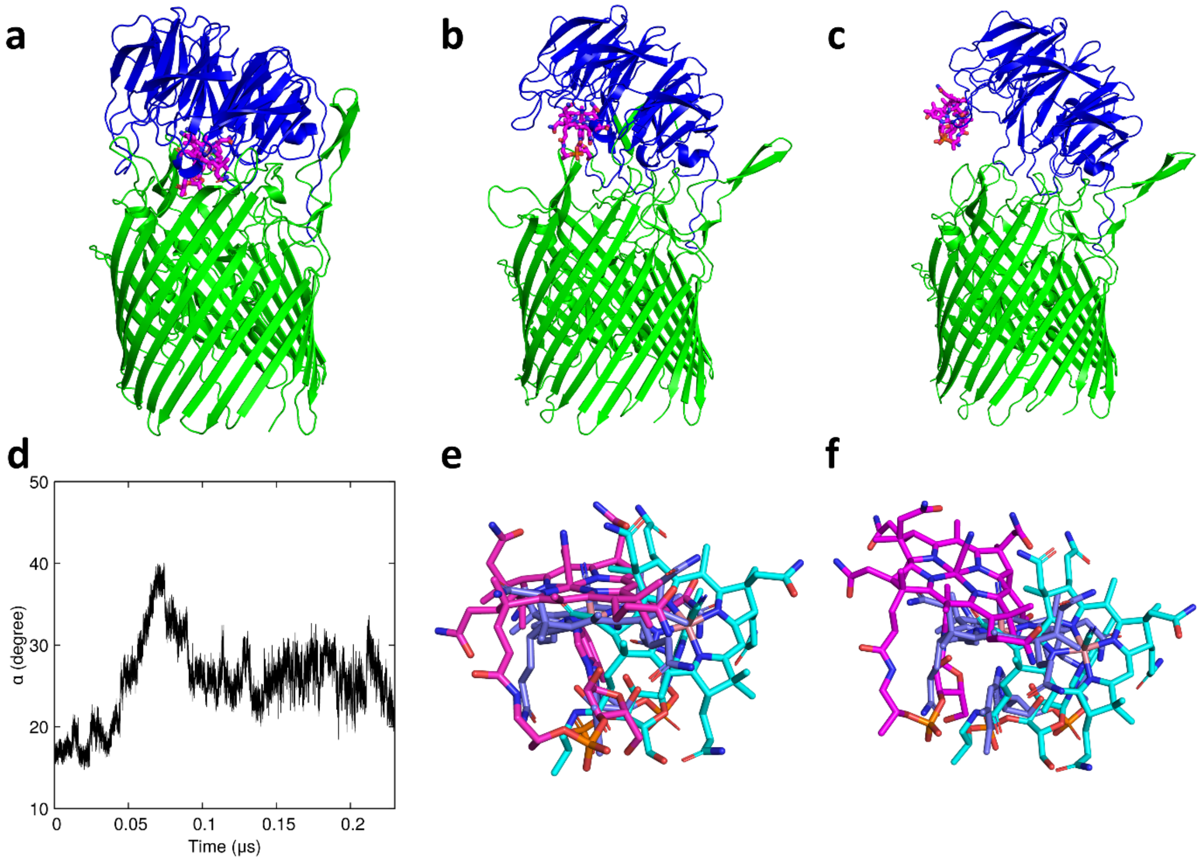


**Supplementary Figure 9 l CNCbl acquisition by BtuB2G2. a**, CNCbl docked into the binding cavity of BtuG2 in the BtuB2G2 complex. **b**, Intermediate structure of BtuB2G2 upon pulling CNCbl along the z-axis, *i.e.*, along the OM surface normal. **c**, Snapshot of the 40° open structure, which was used as input for the unbiased simulation to illustrate the CNCbl acquisition process. **d**, The aperture angle analysis revealed a maximum aperture of ~40° observed at around 75 ns when the CNCbl leaves the protein. **e,f** CNCbl acquisition using additional unbiased MD simulations starting with the structure shown in panel c. The final CNCbl positions in the simulations (magenta) are compared with the docked (cyan; panel a) and BtuG2-CNCbl crystal structure position of CNCbl (light-blue) for simulations 1 (e) and 2 (f) as representative cases.

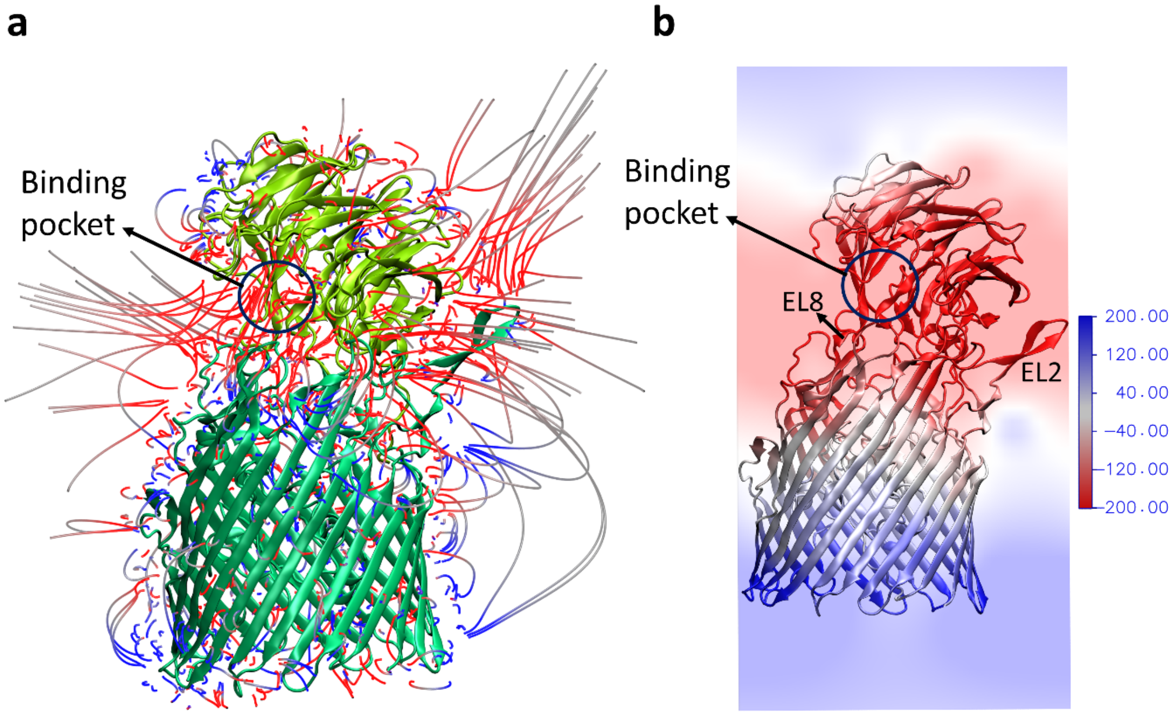


**Supplementary Figure 10 l The electrostatics of the (partially) open BtuB2G2 complex. a,** The electric field line calculation of the BtuB2G2 complex revealed a significant number of electric field lines originating from the binding site of BtuG2 as was observed for the crystal structure of the isolated BtuG2 (see Figure 2e). **b,** The electrostatic potential map of BtuB2G2 demonstrates that the BtuG2 binding pocket in BtuB2G2 has a negative electrostatic potential surface similar to the one for isolated BtuG2 (Supplementary Figure 5c-d).

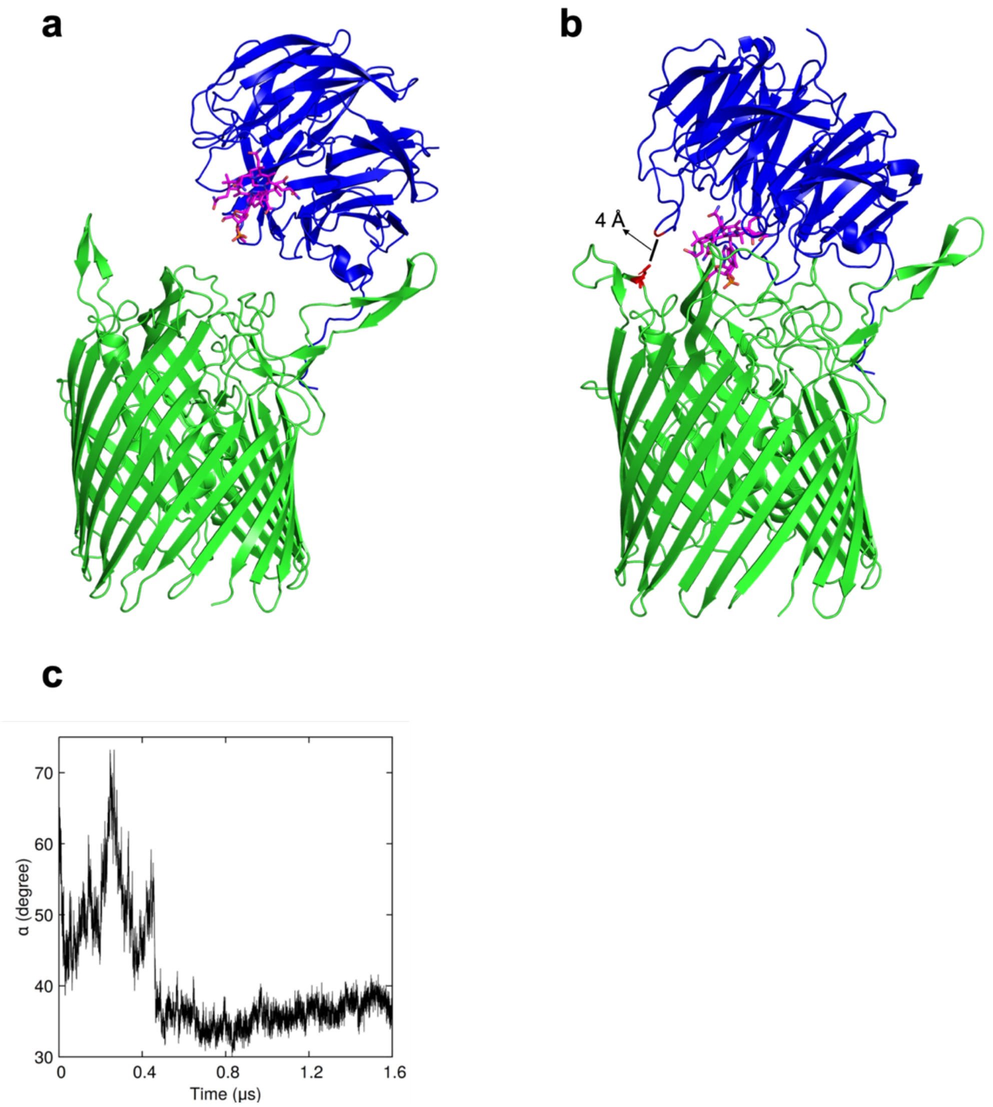


**Supplementary Figure 11 l a, BtuG2 lid closure starting from a wide-open position.** Structural changes of the BtuB2G2 protein during the 1.6 μs unbiased simulation of the CNCbl acquisition process starting from a structure having an aperture of 60° and in which the CNCbl-BtuG2 moiety was replaced by the respective crystal structure. **b**, Partially closed ButG2 lid after 1.5 μs ($\alpha$ = 32-34°). The $\beta$5A loop (residues D236-P249), which hold the CNCbl molecule like pliers, inhibits the closing process. **c**, The distribution of the α angle throughout this simulation. As is evident from panel a, in this wide-open state BtuG2 loses most of its contacts with BtuB2 and the hinge at the back of the complex is no longer apparent, suggesting that this state is unrealistic.

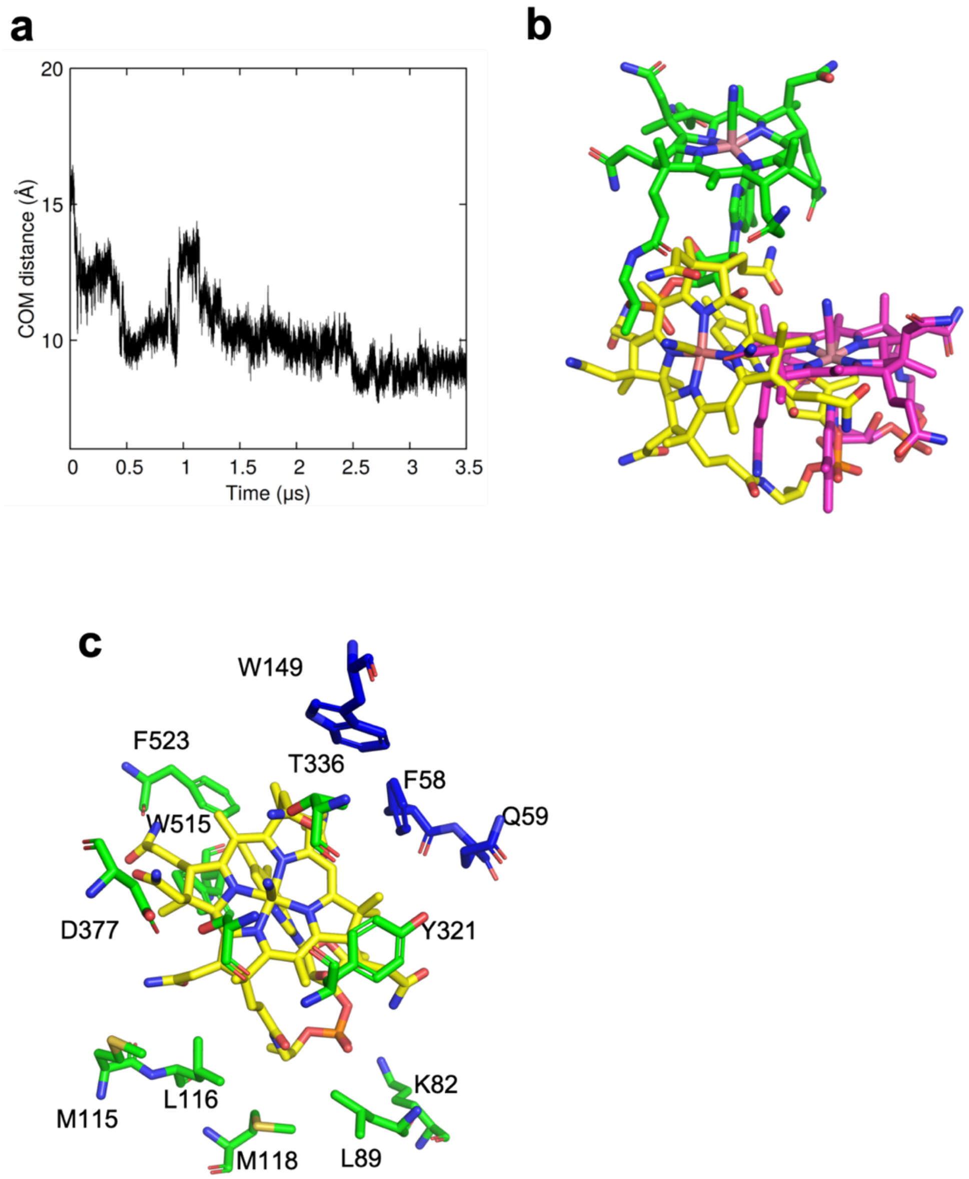


**Supplementary Figure 12 l Translocation of CNCbl from BtuG2 to the BtuB2 binding pocket.** **a,** COM distance between the binding site cavity of BtuB2 and CNCbl. **b**, Snapshot after 3.5 μs, showing how CNCbl moves towards the binding site of BtuB (magenta) from the BtuG2 binding site (green), but becomes stalled close to the BtuB cavity (yellow). The molecule does not move further due to strong noncovalent interactions with residues of BtuB2 and steric hindrance from F523 and W515. **c**, The stalled CNCbl is stabilized by charged and aromatic amino acid residues. The blue amino acids are from BtuG2, and the green amino acids from BtuB2.

**Supplementary Movie 1 l Binding of CNCbl by BtuG2.** Molecular dynamic simulation showing how CNCbl binds BtuG2 in an association-dissociation fashion. β5A loop is represented in red.

**Supplementary Movie 2 l Acquisition of CNCbl by BtuB2G2.** Video showing a 1 μs-long unbiased MD simulation in which BtuG2 closes very slowly, then a CNCbl acquisition event happens and after that further closing of the lid can be seen. β5A loop of BtuG2 is represented in red and EL8 of BtuG are shown in magenta. BtuG2 and BtuB2 are blue and green respectively.
